# The dynamics of oligodendrocyte generation: how distinct is the mouse from the human?

**DOI:** 10.1101/2019.12.23.887174

**Authors:** David G Gonsalvez, Georgina A Craig, Darragh M Walsh, Barry D Hughes, Rhiannon J Wood, Sang Won Yoo, Simon S Murray, Junhua Xiao

## Abstract

Murine oligodendrocyte generation dynamics are considered distinct from those in the human, with implications for cross-species differences in neural homeostasis, injury response and ability to functionally adapt circuits through myelin plasticity. We identify that murine oligodendrocyte precursors do not vary their cell division times *in vivo* and determine how daily production rates change over a lifespan. We show that murine oligodendrogenesis closely resembles what is reported for the human.

## Main

Oligodendroglia comprise a major cellular component of the Central Nervous System (CNS), and are essential for its proper function. Throughout life, oligodendrocyte precursor cells (OPCs) generate the new oligodendroglia that are integrated within neural circuitry. This process can be augmented by neural activity in a form of plasticity known as adaptive myelination^1^. It has been calculated that homeostatic production rates in adult murine white matter are up to 100-fold greater than their human counterparts^2^. Such differences in baseline dynamics may have functional implications in terms of the supply of new oligodendroglia able to partake in myelin plasticity and raises questions as to whether this form of neural adaptation may have a different degree of functional relevance across these species.

Murine OPCs are reported to be a highly proliferative population that maintains a stable growth fraction (GF = cells in G1-S-G2-M, but not G0^3^) with up to 98% in the cell cycle^4–9^. To regulate production OPCs alter their cell cycle length (*Tc*), this can vary from ∼36 hours (h) to ∼864 h depending on age and CNS location^4–9^. However, virtually all of the data on murine OPC proliferation dynamics has been acquired using cumulative labelling with a single thymidine analogue (S-phase tracer)^4–9^. Although this method is widely used to study proliferation in the CNS, the technique can only accurately identify *in vivo* cell cycle parameters if several biological assumptions are satisfied^3^. Unfortunately, parenchymal glial cells violate the key assumptions that underpin these methods (Supplementary Figure 1 **and methods)**. Therefore, reported values for OPC *Tc*, S-phase length (*Ts*) and GF may not appropriately reflect *in vivo* dynamics. Furthermore, modelling that uses these data may not accurately estimate the rates of murine oligodendrocyte production, and any subsequent conclusions about cross species developmental differences could be inaccurate.

**Figure 1.**
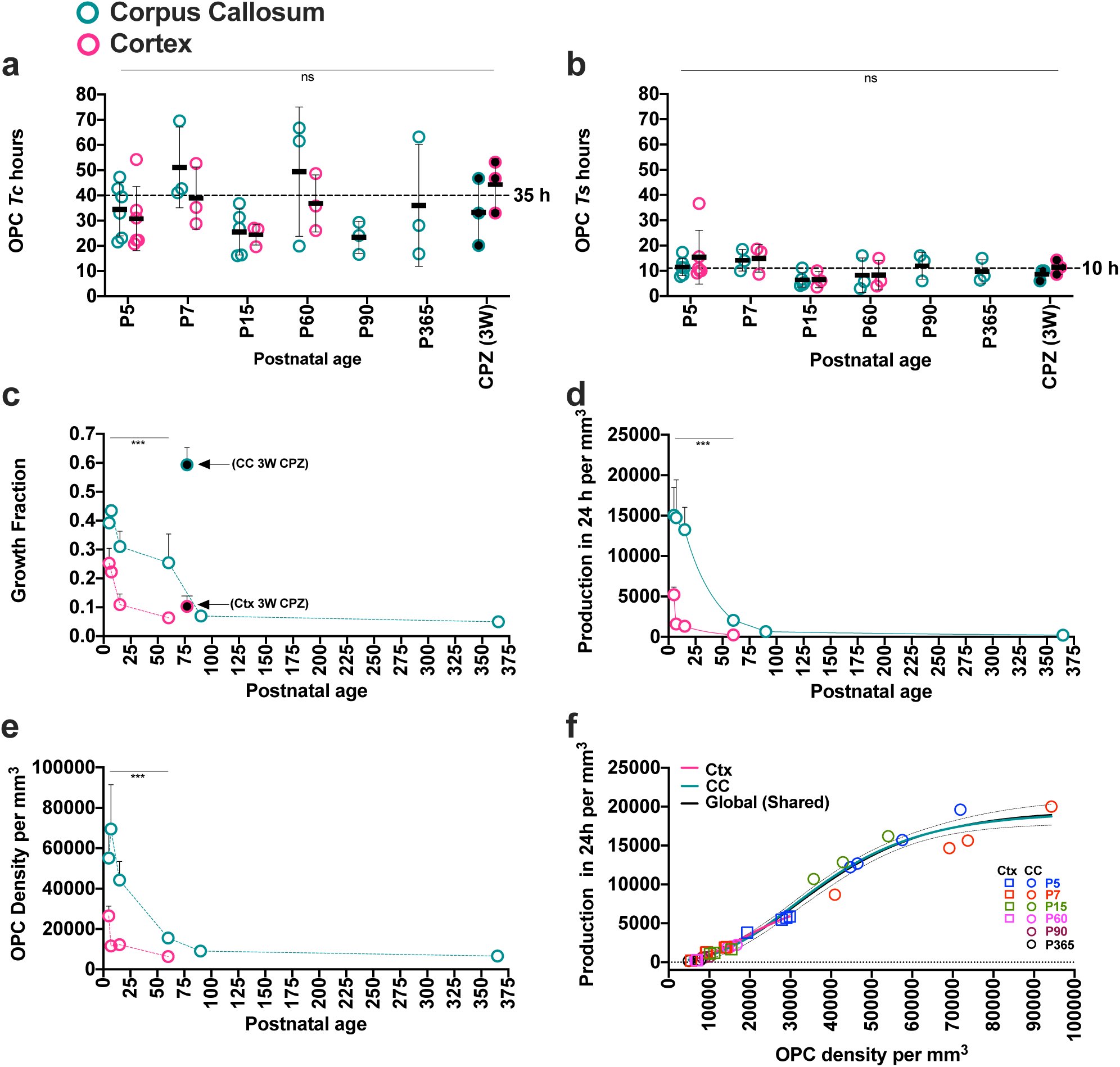
OPC production rates decline abruptly in early post-natal development. **a)** OPC cell cycle length (*Tc –* hours) throughout life in the corpus callosum and cortex. In all figures filled circles represent data from individual mice feed 0.2% cuprizone for 3 weeks and collected at P77 (3W CPZ). **b)** OPC S-phase length (*Ts* – hours) through life in the corpus callosum and cortex. **c)** OPC Growth Fraction (GF) throughout live in the corpus callosum and cortex. Callosal OPC GF is dramatically elevated in response to cuprizone-induced demyelination in the corpus callosum (3W CPZ arrows). The GF was determined using Ki67 immunoreactivity (Supplementary Figure 3 **and 4**). **d)** Daily OPC production rates in the corpus callosum and cortex during postnatal development. (**see** Supplementary Figure 9a **for statistical comparisons between ages**). **e)** Developmental changes in the volumetric density of OPC in the corpus callosum and the cortex (**for additional statistics see** supplementary figure 10f). **f)** The volumetric density of OPC is positively related to their production rate. The density to production relationships for the cortex (orange line) and corpus callosum (blue line) were not significantly distinct and followed a global relationship (black line - **see** Supplementary Figure 9b **for the equations and R^2^ values).** Statistics, at each timepoint minimum n=3, 2-way ANOVA interaction significance indicated by *** where P < 0.001. In A – E, error bars = SD. In F, the broken lines indicate 95% confidence intervals.

A double S-phase labelling approach incorporating Ki67 immunoreactivity overcomes the technical limitations of single S-phase tracer cumulative labelling (Supplementary Figures 2 **& 3)**^10^. Using this approach, we identify that the average OPC *Tc* is 35 h ± 14 h SD and remains remarkably stable irrespective of age or CNS location (Figure 1A). This range of OPC *Tc* values is concordant with the range of *Tc* values reported for rodent OPCs *in vitro* ^11^. We also found that the average OPC *Ts* was 10 h (± 5 h SD) and did not significantly vary across age or location (Figure 1B). In contrast, OPC GF decreased dramatically with age and was significantly distinct between the corpus callosum and cortex at all time points assessed (Figure 1C **and** Supplementary Figure 3). Importantly, the developmental changes we identified in the murine callosal GF via Ki67 immunoreactivity resemble what has been reported for the human corpus callosum (Supplementary Figure 4). Moreover, the GF values we identified are consistent with the fraction of OPCs expressing cell proliferation genes as determined by single cell RNA sequencing at comparable ages^12^. In the context of demyelinating injury, we observed no change to OPC *Tc* or *Ts* (Figure 1 **A and** B) in response to 3 weeks of the cuprizone-induced CNS demyelination (see methods). That said, cuprizone led to a dramatic increase in callosal OPC GF (Figure 1C **and** Supplementary Figures 3 & 5). Collectively, these data provide evidence that OPCs alter their proliferative behaviour principally through changes in their GF, while maintaining a consistent *Tc* and *Ts in vivo*.

**Figure 2.**
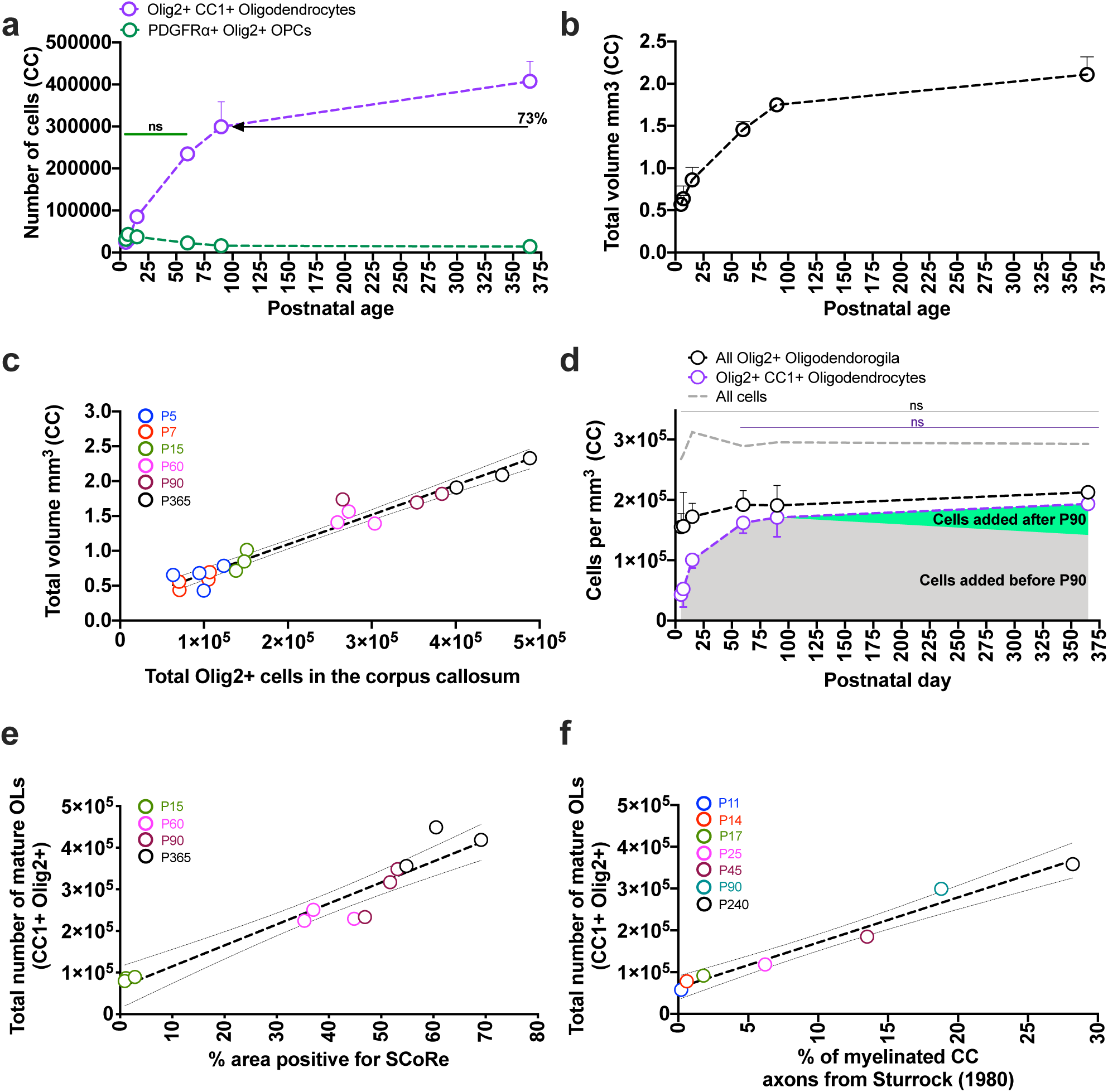
The majority of mature callosal oligodendrogila are generated within the first 10% of the murine lifespan. **a)** The total number of mature oligodendroglia and OPCs in the corpus callosum over the lifespan. By P90, 73% of the total number of mature oligodendroglia observed at P365 have been generated. There is no significant change in total OPC number from P5 to P60. Beyond P60, the total pool of OPCs is depleted gradually over the course of development (for additional statistics see Supplementary Figure 7). **b)** The volume of corpus callosum, bound by the cingulate bundles laterally, the genu rostrally and selenium caudally, increases with age (also see supplementary Figure 3). **c)** The total number of oligodendroglia (all Olig2+ cells) has a strong positive correlation with the total volume in the corpus callosum. Each symbol represents data form an individual animal. Pearson’s r = 0.98, R2 = 0.95 and P< 0.0001 (error = 95% CI). **d)** Age-related changes in cell density within the murine corpus callosum. The density of mature oligodendroglia increases until P60, after which the increase in number mature callosal OLs occurs with no change in volumetric density (see additional statistics see Supplementary Figure 8). **e)** The total number of mature oligodendroglia (CC1+ Olig2 positive cells) is strongly correlated with the increase in % of callosal area positive for SCoRe. Each symbol represents data form an individual animal. Pearson’s r = 0.95, R2 = 0.91 and P< 0.0001 (error = 95% CI). **F)** The number of mature oligodendroglia is strongly correlated with the % of myelinated axons determined by Sturrock (1980)16. Each symbol represents mean values for total CC1+ Olig2 positive cells where at minimum n=3/age and mean values of the % of myelinated axons from Sturrock (1980 – table 1)16. Pearson’s r = 0.99, R2 = 0.97 and P< 0.0001 (error = 95% CI).

**Figure 3:**
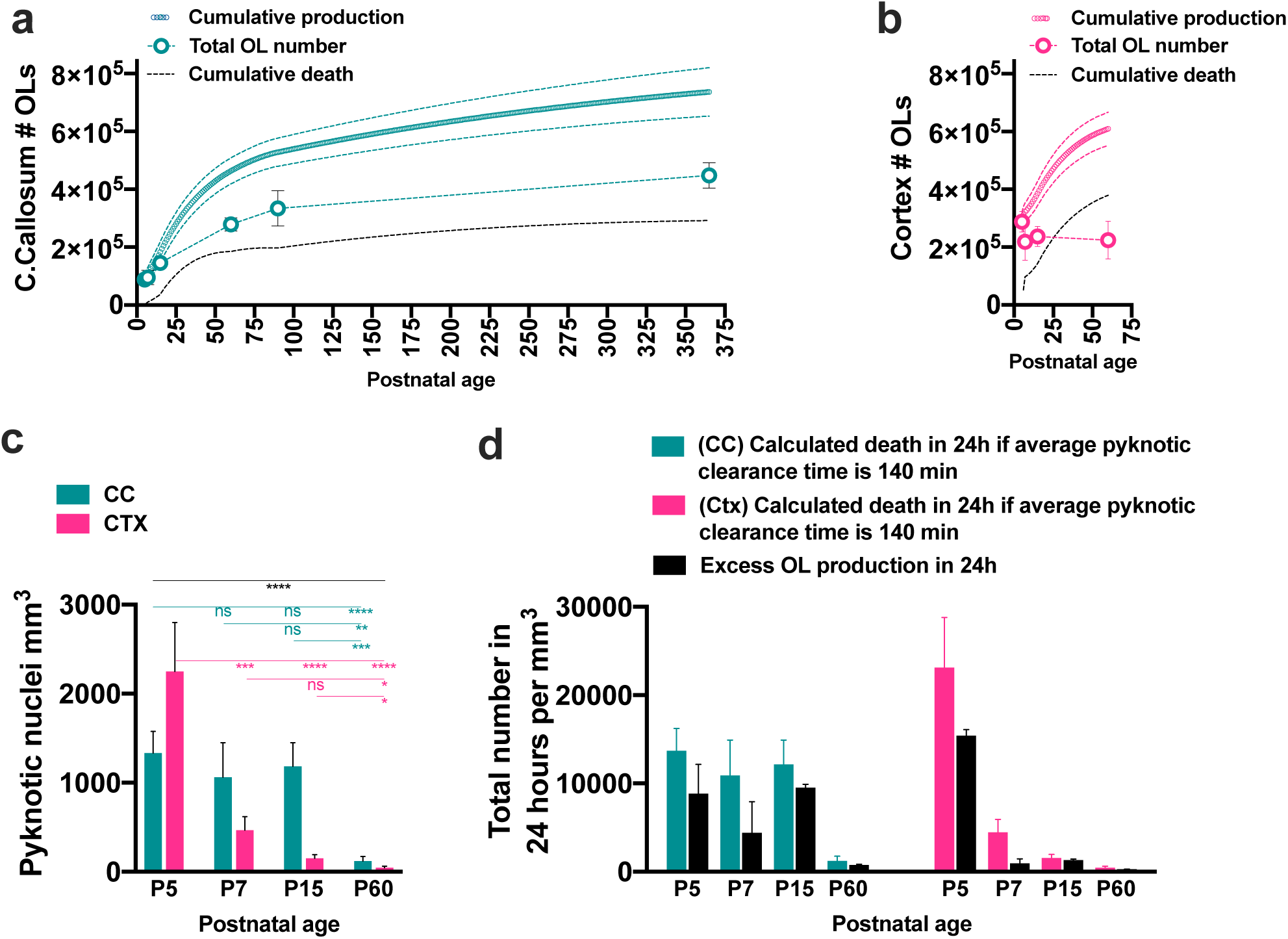
OPCs generate oligodendroglia in excess thought development. **a)** Cumulative daily production by OPCs in the corpus callosum (teal circles ± SD) exceeds the change in total oligodendrocyte number as determined by stereology (bold teal circles ± SD). The difference between production and the increase in total oligodendroglial number represents cell death in the system (black broken line). **b)** Cumulative daily production by OPCs in the cortex (pink circles ± SD), exceeds the change in total oligodendroglial number determined by stereology (bold pink circles ± SD) and cell death (black broken line). **c)** Spatiotemporal distinctions in the density of pyknotic nuclei between the corpus callosum and cortex. **d)** Total pyknotic nuclei counts were used in conjunction with published data on the average time it takes for a dead cell to be cleared from the CNS (140 min)^19^ to calculate total daily cell death per mm^3^ (see methods). This was compared to the values for daily excess cell production identified in A and B (these values are represented by the solid black bars). The calculated total daily death, using pyknotic nuclei counts and estimated clearance times for dying cells in the cortex, exceeded the level of daily excess OPC production identified for that equivalent day. Statistics, at each timepoint minimum n=3, 2-way ANOVA analysis with multiple comparisons significance: *, **, ***, **** = P<0.05, P<0.01, 0.001, and p<0.0001 respectively.

We next used design-based stereology to quantify the developmental changes in tissue volume (Supplementary Figure 6) and absolute numbers of oligodendroglia within an anatomically defined region of the corpus callosum (Supplementary Figure 7) and cerebral cortex (Supplementary Figure 8). Combining the stereological counts with our cell cycle data, it was possible to identify how daily OPC production rates change with age. Per unit of volume, callosal and cortical OPC production rates decline sharply during early postnatal development (Figure 1D **and** Supplementary Figure 9A). Regional distinctions in how OPC density (Figure 1E **and** Supplementary Figure 10F) and population GF (Figure 1C) change underpin the striking differences in the production rates between the two CNS regions (Figure 1D). We next plotted OPC production against density and this revealed a striking relationship – the number of new OPCs produced for a given density was consistent regardless of age or CNS location during normal development (Figure 1F **and** Supplementary Figure 9B). We find that OPCs maintain an elevated GF when at a higher volumetric density under normal homeostatic conditions (Supplementary Figure 9C). These findings have been validated using data independently collected by Hughes et al^13^ **(data plotted onto** Supplementary Figure 9B and C).

In the human corpus callosum the density of proliferating OPCs rapidly declines in early development and 80% of the total number of mature adult oligodendrocytes are generated by 5-10 years of life^2^. Consistent with these findings, we found that 73% of the total number of mature callosal oligodendroglia at postnatal day 365 (P365) have been generated by P90 (Figure 2A). If we consider relative differences in lifespan (∼90 years for a human^2^ and ∼900 days for a laboratory mouse^14, 15^) and the very low rates of adult callosal OPC production (Figure 1D), the majority of callosal oligodendrogenesis is complete within the first 10% of the lifespan in both species. We also show that the total number of mature oligodendroglia increased until P365 (Figure 2A **and** Supplementary Figure 7). Interestingly, total callosal volume increases to accommodate the rise in total oligodendrocyte number (Figure 2B). We plotted the total number of oligodendroglia against callosal volume and revealed a very strong positive linear relationship **–** the callosum maintains a stable cellular density as it grows in development (Figure 2C **and** D). This means using density measures alone it may not be possible to accurately capture how populations change within the oligodendrocyte lineage during development. For example, although we identified a dramatic decline in the volumetric density of callosal OPCs from P5 to P60 (Figure 1E **and** Supplementary Figure 10F), there was actually no significant change in their total number over this period (Figure 2A **and** Supplementary Figure 7). The changes in callosal OPC density are not due to an enhanced rate of differentiation that depletes the total number of OPCs. Instead, high rates of cellular production and subsequent integration expands the total number of mature callosal oligodendroglia and total callosal volume (Figure 2 A **and** B), and this is what causes the fall in OPC volumetric density **(compare** Figures 1E and 2A). Beyond P60, total callosal OPC number gradually declines with age (Figure 2A **and** Supplementary Figure 6), suggesting over time the rate of proliferation is exceeded by the rate of differentiation within this self-renewing pool of cells.

We next plotted the total number of mature callosal oligodendrocytes against myelin, assessed by spectral confocal reflectance microscopy (SCoRe), and found a strong positive linear relationship between the area positive for SCoRe and the number of mature oligodendrocytes in the corpus callosum (Figure 2E **and** Supplementary Figure 11). To validate this, we plotted the total number of mature oligodendrocytes against an independent report assessing the proportion of myelinated axons in the corpus callosum assessed by Transmission Electron Microscopy^16^, and again found a strong positive linear relationship (Figure 2F). Our data does not exclude the possibility that existing cells may re-model the complement of myelin they produce, however it argues that increasing the total axonal area covered by myelin depends on the addition of new myelinating oligodendroglia. This finding is consistent with recent reports detailing how myelination occurs in the adult murine cortex^14, 15^.

After 5-10 years of age in the human, the number of newly generated oligodendroglia that become integrated into the corpus callosum each year accounts for 0.33% of the total mature oligodendroglia in the tissue^2^. After P90 in the murine corpus callosum, newly generated mature oligodendroglia expands the existing number of these cells by a rate of 0.44% every 10 days (see methods for calculation). Accounting for the relative differences in lifespan between these species, these rates are not strikingly distinct. However, it is important to point out that in our case the 0.44% rate represents only the addition of new mature oligodendroglia and does not include any cellular replacement or turnover. We cannot exclude the possibility that existing mature callosal oligodendroglia may die and subsequently be replaced while their total number expands. However, our data finds that callosal OPC production rates are extremely low beyond P90 (Figure 1D). This, along with compelling evidence that mature murine oligodendrocytes are remarkably stable^14, 15, 17^, makes it difficult for us to envisage a situation where a high rate of cellular turnover occurs as part of normal homeostasis in the murine corpus callosum.

We know that programmed cell death eliminates a proportion of the nascent pre-myelinating oligodendroglia and represents a mechanism that can limit the rate of cellular integration^18^. Using the daily OPC production rates and stereological counts, we were able to extrapolate total cumulative production and compare this against the actual change in total oligodendroglial number in corpus callosum (Figure 3A) and cortex (Figure 3B). In both regions, cumulative daily OPC production exceeded the increase in absolute number of all oligodendroglia as determined by stereology (Figures 3A **&** B). The only way we could account for the excess OPC production was to assign this as cell death (see methods). To validate these data, we quantified developmental changes in pyknotic nuclei (Figure 3C **and** Supplementary Figure 12). Combining the pyknotic nuclei counts and data on the average clearance times for dead cells in the developing CNS^19^, it was possible to estimate total daily death at different times in the tissue (Figure 3D). We found that at all ages our values for excess production (cell death) were within the estimates for total cell death (Figure 3D). In addition to this, sensitivity analysis was performed using a fixed OPC *Tc* value of 35 hours, this generated a cumulative production curve nearly identical to what we observed when using the measured *Tc* values (Supplementary Figure 13). However, a systematic over/underestimation of OPC *Tc* by ± just one standard deviation had a profound influence total cumulative production (Supplementary Figure 13).

There is strong evidence that changes in neural activity can augment the homeostatic rates of OPC production and new oligodendrocyte integration *in vivo*^1, 20^. Furthermore, blocking the rapid integration of new oligodendroglia abrogates particular behavioural adaptions that normally occur in response to altered experience^21^. Our data demonstrate that the volumetric density of OPCs and their production rates decline in a regionally-specific manner with age (Figure 1D **and** E). These spatiotemporal changes will influence the probability that an OPC, or newly generated oligodendrocyte, will be in close proximity to an axon whose activity has been altered. In early postnatal life, the density of OPCs and their production rates are dramatically distinct when comparing between the corpus callosum and cortex, however these regional differences become much less pronounced with age (Figure 1D **and** E). These findings may help to explain why the same level of increased neural activity elicits profound regional distinctions in OPC behaviour in young mice, but a diminished and more uniform response across the same CNS regions in older animals^1^.

Collectively our data provides new fundamental insights on how OPCs function to regulate cell production *in vivo*. We demonstrate that like other self-renewing cell populations within the CNS, oligodendrocyte generation dynamics are largely conserved between the mouse and human.

## Methods

### Animals

C57BL/6 mice were used in all experiments. Mice were housed in specific pathogen-free conditions at the Melbourne Brain Centre Animal Facility. All animal procedures were approved by The Florey Institute for Neuroscience and Mental Health Animal Ethics Committee and followed the Australian Code of Practice for the Care and Use of Animals for Scientific Purposes.

### Tissue collection

Prior to tissue collection all mice were perfused using phosphate buffered saline (PBS) followed by 4% paraformaldehyde (PFA). Brains were dissected and tissue blocks prepared using the appropriate coronal brain matrix (Harvard Apparatus). All brains were post-fixed overnight in 4% PFA, then washed in PBS and transferred to a 30% sucrose solution. Following this, the tissue blocks were imbedded in OCT and snap-frozen using iso-pentane cooled by dry ice. In all cases blocks were serially sectioned caudally to rostrally with all sections carefully accounted for. Sections were cut at 25µm.

### Glia violate the assumptions required for single thymidine-analogue cumulative labelling

Thymidine-analogue cumulative labelling is an assay designed to identify cell cycle parameters of dividing cells *in vivo*^3^. The method involves continuous delivery of a thymidine analogue (S-phase tracer) that is incorporated into DNA of cells as they progress though the S-phase of the cell cycle (Supplementary Figure 1). This enables one to measure the rate at which a population of dividing cells becomes saturated with the S-phase tracer, along with the absolute fraction of cells that becomes saturated within the population (Supplementary Figure 1C). Using this data, it is possible to determine key cell cycle parameters^3^. However, one caveat to using thymidine analogues is that once incorporated into the DNA of a cell, the cell is permanently labelled. Therefore, specific criteria are required in order to exclude any labelled cell that becomes quiescent or moves into G0 over the labelling period and the method was purpose-designed for use in tissues in which proliferating cells are anatomically separated from cells that exit the cell cycle or move into G0^3^. For example, in the dentate gyrus of the hippocampus, it is assumed that any daughter cell that exits the cell cycle and moves into G0 will leave the anatomically-defined proliferative area, the sub granular zone (SGZ), by migrating away into the granule cell layer (GCL)^3^ (Supplementary Figure 1B). The labelled cells that migrate out of the proliferative zone are purposefully not included in the counts that generate the cumulative plots (Supplementary Figure 1B). Formally, the experimental design for single thymidine- analogue S-phase cumulative labelling requires that the following assumptions are satisified3:

1) the cell population is growing at a steady-state, i.e., one-half of the daughter cells leave the proliferative population and one-half remain in the proliferative population;
2) no cells within anatomically-defined proliferative zones can exit and re-enter the cell cycle (move into and out of G0); and
3) cells within the population divide asynchronously.

OPCs do not satisfy these assumptions. For parenchymal glia, there is no anatomical separation between labelled daughter OPCs that exit the cell cycle and move into G0 compared to cells that remain in cell cycle (Supplementary Figure 1D & E). Therefore, it is impossible to segregate a labelled OPC that becomes quiescent and moves into G0, from cells that are actively dividing when using a single S-phase tracer alone. This means that the cumulative labelling method cannot accurately report on the true Growth Fraction within the population. Instead, the population GF will be overestimated in these circumstances (Supplementary Figure 1 E–F). This is reflected in experimental evidence that near-complete saturation of all OPCs occurs after long enough exposure to a non-toxic level of S-phase tracer^8^. Furthermore, the rate at which the pool of OPCs becomes saturated will be influenced by their rates of differentiation and cell death, in addition to the proliferative behaviour of the OPCs. This means that for any OPC cumulative curve plot, the measured saturation time is not indicative of cell cycle dynamics alone and cannot be reliably used to accurately interpret the S-phase length or cell cycle length for OPCs.

### Identifying cell cycle parameters using double S-phase labelling with Ki67

To overcome limitations with single S-phase cumulative labelling, we previously devised an approach that combines an acute double S-phase labelling approach^22^ with Ki67 immunoreactivity^10^. This enables identification of key cell cycle parameters in cell populations where there is no anatomical separation between actively dividing cells and those in G0^3^. Using this approach, the following parameters can be identified:

*<u>The S-phase length (Ts)</u>* – is the time a cell spends in the S-phase of the cell cycle. This is determined using the double s-phase injection protocol^10, 22^. Mice were injected intraperitoneally, initially with BrdU (Roche Diagnostics) at 100 µg.g^-1^ body weight and 2 h later with EdU (Invitrogen) at 50 µg.g^-1^ body weight. Thirty minutes after the injection of EdU, mice were sacrificed by lethal injection of pentobarbitone and perfused. Identification of the length of s-phase can be determined using the following relationship:

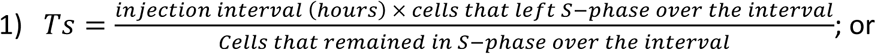

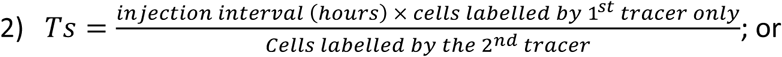

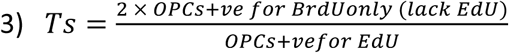

The 2-hour interval between injections of BrdU and EdU was selected based on previous literature assessing cell cycle dynamics for neuroblasts and glial cells in neural tissues^10, 22^. This technique requires accurate identification of the proportion of cells that left S-phase over the injection interval^22^. The order of injection BrdU first, followed second by EdU is very important. A carbon alkyne is engineered into the EdU and the azide-N_3_-fluorophore used to identify the EdU is attached via a Copper-catalyzed azide-alkyne cycloaddition. This means it is not possible for the EdU fluorophore to cross-react or falsely detect the BrdU in the system (Supplementary Figure 2). The double S-phsae labelling protocol requires that a measurable number of cells pass through S-phase over the injection interval^22^. Unfortunately, in the cortex at P90 and P365, too few cells passed though S-phase over the injection interval to reliably report on the S-phase length, so cortical assessments were performed until P60. We were able to obtain S-phase lengths for animals at P365 only in the corpus callosum. However, it must be noted that we observed very few OPCs positive for BrdU only at P365.

*<u>The Instantaneous Labelling Index (Li_0_)</u>* – is the fraction of cells actively in S-phase at any instant. The double s-phase labelling technique used to identify *Ts* enables identification of the fraction of OPCs labelled by EdU (a 30min pulse) and BrdU (a 150 min pulse) in each animal. The LI_0_ is the y-intercept from a least of squares linear equation derived by passing a line though measured fractions of labelled OPCs at 30 min and 150 min of exposure time^10, 22^ (Supplementary figure 2).

*<u>The Growth Fraction (GF)</u> –* is the fraction of cells within a population that are actively in the cell cycle, or all cells excluding those that are in G0^3^. This was determined using immunofluorescence for Ki67 and PDGFRα (Supplementary Figure 3)^10^. Ki67 expression discriminates cells in G1-S-G2-M from cells in G0^10, 23, 24^. The Ki67 protein has a short half-life (60-90min) and its levels change in different phases of the cell cycle^24–26^. Although Ki67 is routinely used as a binary marker of proliferating cells, there is evidence that under some *in vitro* conditions a lag in the onset of Ki67 expression may occur early in G1^24–26^. While this lag is not long (1 −2 hours) and is only observed when cells have been deprived of growth factors then re-introduced to mitogens, or when cells are maintained in G0 by experimental manipulation of mitotic checkpoints and then returned into the cell cycle, nevertheless it is possible that a lag in onset of Ki67 expression may occur during early G1 for *in vivo* OPCs that have been quiescent then re-enter the cell cycle^24–26^. To test if OPCs *in vivo* may have a significant lag in the onset of Ki67 expression we plotted the GF/LI_0_ (The fraction of dividing cells, identified by Ki67 / fraction of cells in S-phase identified by double S-phase labelling). If quiescent OPCs that move from G0 to G1 have some significant lag in the onset of Ki67 expression, this must lead to a decrease in the GF/LI_0_ fraction as OPCs become less proliferative with age. We found no significant change in the GF/LI_0_ fraction across age, location or in response to cuprizone (Supplementary Figure 2D). Our data does not support a hypothesis that an extended lag in the onset of Ki67 occurs during G1 phase of the cell cycle of OPCs *in vivo*. High resolution images were assessed in all cases for determining the GF. PDGFR⍺+ pericytes were always excluded from counts, this was done based on their morphology and proximity to parenchymal blood vessels^27^ (Supplementary Figure 3B).

*<u>The cell cycle length (Tc)</u> –* is the time taken for a cell to complete cell division, or the time taken to pass through G1, S, G2 and M. This can be determined using the formula^3^:

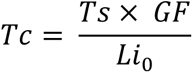

One benefit of our approach is that all of the cell cycle parameters Ts, LI_0_, GF and Tc can be determined for individual animals, rather than relying on pooled data. This is not possible using single S-phase cumulative labelling, where pooled averages from multiple animals are required to determine each single parameter at a given age.

### Immunohistochemistry

We devised a sampling strategy to obtain cell cycle data and stereological counts from the same individual, with sections processed in one of 4 ways:

1) Sections were processed for PDGFRα immunofluorescence (Table 1) before being treated with 2N HCl for 30 min at room temperature and then 0.1 M sodium tetraborate. They were then processed for BrdU immunofluorescence and finally reacted for EdU as per the manufacturer’s instructions for the Clik-iT EdU Alexa Fluor 647 imaging kit (Supplementary Figure 2);
2) Sections were subjected to antigen retrieval (10 min at 90°C in 0.01 M citrate buffer, pH 6.0), reveal then processed for Ki67 and PDGFRα immunofluorescence (Table 1) 27 (Supplementary Figure 3 and 5);
3) Sections used for stereology counts were processed for Olig2, PDGFRα and CC1 immunoreactivity (Table 1), this enabled identification of phenotypically distinct cells within the oligodendrocyte lineage (Supplementary Figure 6); or
4) Sections were processed for IBA-1 immunoreactivity (table 1) in antisera free of any detergents that enabled combined SCoRe assessments in cuprizone fed mice^28^ (Supplementary Figure 5).

| *Antibodies and reagents* | *Source* | *Concentration* | *Reference* |
| --- | --- | --- | --- |
| Goat anti Platelet derived growth factor alpha (PDGFRα) | AF1062, R&D Systems | 1:200 | Fletcher et al (2018) |
| Mouse anti-APC (AB-) MAB CC1 | OP80, CalBioChem | 1:200 | as above |
| Rabbit anti Olig2 - Rabbit | AB9610, Millipore | 1:200 | as above |
| Rat anti 5-Bromo-2'-deoxyuridine (BRdU) - Rat | Abcam (ab6326) | 1:40 | Gonsalvez et al (2013) |
| Rabbit anti Ki76 -Rabbit | RM-9160, Thermo | 1:100 | as above |
| 5-Ethyl-2-deoxyuridine 647 Click reaction kit | Life Technologies | as per kit | as above |
| Bisbenzamide - Hoechst 33342 | Sigma | 1 μg /ml | as above |
| Goat anti IBA-1 | Abcam (ab5067) | 1:200 |  |

### Cuprizone induced demyelination

8-week-old mice were fed 0.2% cuprizone in normal chow (Teklad Custom Research Diets)^28^. After 3 weeks of dietary cuprizone supplementation mice were sacrificed and tissue processed as described previously. Animals were injected and all tissue was processed as previously described. Confirmation that cuprizone had elicited inflammation and demyelination was determined using SCoRe imaging and immunofluorescence for IBA-1+ microglia (Supplementary Figure 5)^28^.

### Imaging, Spectral Confocal Reflectance Microscopy and image analysis

All images were captured on either a Zeiss LSM 880 (Airyscan), Zeiss LSM780 confocal microscope, or Zeiss Axio M2 with apotome. Spectral Confocal Reflectance Microscopy (SCoRe) was performed using either the Zeiss LSM 880 or LSM780 microscopes. Both microscopes were fitted with a water immersion objective (Zeiss W Plan-Apochromat 40 /1.0 NA DIC) using 458, 561, and 633-nm laser wavelengths passed through the Acousto-Optical Tunable Filters (AOTF) 488–640 filter/splitter and a 20/80 partially reflective mirror. The reflected light was collected using three photodetectors set to collect light through narrow bands defined by prism and mirror sliders, cantered around the laser wavelengths 488, 561, and 633 nm. The channels from each photodetector were then overlaid as one composite image for analysis. In all cases SCoRe images were pseudo coloured in cyan. All images were acquired using the same process. The tile scans captured for each section were taken in a single z-plane at a minimum at a depth of 3 micron from the surface. Only compact myelin reflects light to produce a positive signal. Positive pixels were identified on a minimum threshold cut-off using the threshold function in Fiji. Measurements of the resulting area were divided by the total area of the ROI^28, 29^. A minimum of n=3 was captured for each age.

### Design-based Stereology

The anatomical structures of the cingulate bundles laterally, and the rostral and caudal midline unions of the corpus callosum were used to define the anatomical boundaries of the cortical and callosal tissue sampled (Supplementary Figure 6). To ensure systematic random sampling, brains were serially sectioned (25μm thick). This approach meant any developmental changes in the tissue volumes bound by these anatomical landmarks could be determined (Supplementary figure 6). All nucleated cells in the corpus callosum and cortex were counted using the optical fractionator method with Stereo Investigator version 11.01.02; (MBF Bioscience). The total section thickness was measured at every single counting site to maximise the accuracy in volume estimates. To identify the total number of all oligodendroglia, OPCs, oligodendrocytes and non-oligodendrocytes in each animal, sections were also processed for multi-label immunofluorescence (Supplementary Figure 6). The grid frame and counting window sampling protocol used ensured a Gundersen M0 Coefficient of Error (M0 CE) of <0.05, significantly less than the commonly accepted M0 CE variance value of 0.1. Total cell numbers for the corpus callosum and cortex (Supplementary Figure 7 and Supplementary 8 respectively). Total cell density per mm^3^ for the corpus callosum and cortex (Supplementary Figure 11)

### Calculating daily OPC production

Daily production was calculated by identifying the total numbers of dividing OPCs. This was achieved by multiplying the total number of OPCs identified though stereology (Supplementary Figure 7-8), by the Growth Fraction identified by Ki67 immunoreactivity (Figure 1C). It was then possible to calculate daily cell production in the following way:

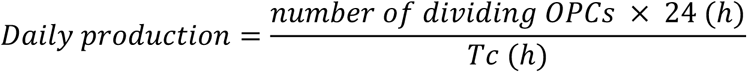

The stereological approach included measuring the total tissue volume (Supplementary Figure 6). It was therefore possible to convert total daily production to a volumetric density:

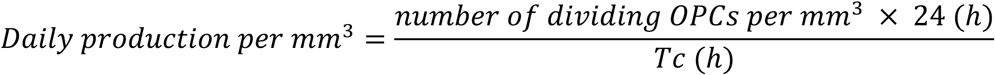

For the corpus callosum, we determined cell cycle parameters and stereological data at the following data points: P5, P7, P15, P60 and P90 and P365. To determine the number of OPCs, linear interpolation between the measured mean values of the GF and Tc was used, enabling us to model how production changed between the measured data points.

### Calculating the % expansion rate for total mature oligodendroglia in the corpus callosum

To estimate the % expansion rate for mature oligodendroglia in the murine corpus callosum that occurs after P90, we make the following assumptions:

1) That beyond P365 there is no significant increase in total mature callosal oligodendroglia; &
2) The average murine lifespan is 900 days

To determine the % rate of expansion we performed a simple calculation:

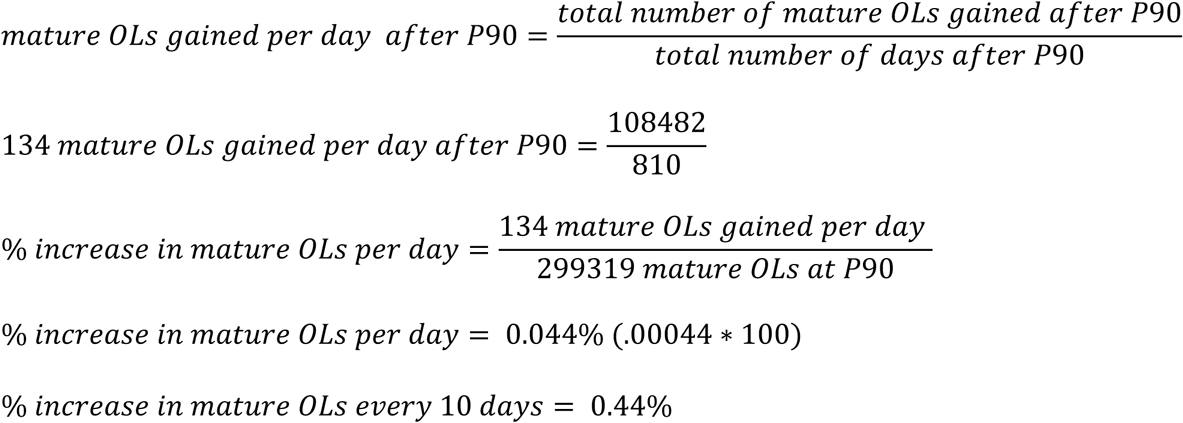

### Non-linear regression analysis

We plotted cortical and callosal OPC volumetric density against daily production (Figure 1E). This did not follow a linear relationship, but we found that the data fitted the Gompertz non-linear regression model:

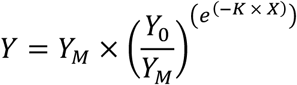

where *X* is the OPC density per mm^3^ and is the daily production per mm^3^. Two of the three parameters *Y*_0_, *Y*_M_ and *K* have simple interpretations: *Y*_M_ is the extrapolated production rate at mathematically infinite OPC density and serves as an estimate of the production rate at the highest achievable OPC density, while *K* (its units being the inverse of those of *X*) controls the rate of variation of production with OPC density. The parameter *Y*_0_ can be interpreted mathematically, but only with some diffidence, as the production rate extrapolated to zero OPC density. The fitting was performed with GraphPad Prism 8 for MacOS (see www.graphpad.com/guides/prism/8/curve-fitting/reg_gompertz-growth.htm, accessed June 19, 2019, but note that their interpretation of 1⁄*K* as the *X* value of the inflection point is not correct, the true inflection point being located at ln{ln[*Y*_M_/*Y*_0_]}/*K*). The fit was confirmed using the NonlinearModelFit function in Mathematica Version 11 (Wolfram Research), with the Prism 8 fitting parameters rounded to two significant figures as initial values in the parameter search and all parameters constrained to be positive. The values for *Y_M_*, *Y*_0_, *K* and the *R*^2^ for each equation are given in (Supplementary Figure 9 B).

To identify if OPC volumetric density was correlated with their probability for division we plotted OPC density against OPC GF (Supplementary Figure 9 C). We fit the data using the Padé (1,1) non-linear regression model in GraphPad Prism 8:

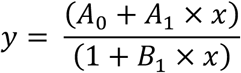

A comparison of fits with the null hypothesis that one curve fits all data points was not possible to reject P = 0.3715. This data indicates that irrespective of age or location, OPCs GF was positively correlated to their volumetric density under normal homeostatic conditions in vivo (Supplementary Figure 9C).

### Modelling cumulative production and death

Cumulative production was identified by cumulatively adding daily production values to the initial number of total OLs (All Olig2+ cells) determined by stereology at P5. Cumulative production exceeded the increase in total OL number determined by stereology in both the corpus callosum and cortex (Figure 3 A–B). The differences between the cumulative curve, and the measured number of OPCs in the tissue represents the excess cell production (Figure 3 A–B). The only way that we could account for the excess OPC production was to assign this as the cell death in the system. This is based on the following assumptions:

1) PDGFRα+ Olig2 + OPCs only produce cells that contribute to the oligodendrocyte lineage^8^;
2) Within the large anatomically-defined volumes of cortex and corpus callosum assessed, net OPC migration = 0. OPCs are highly mobile and although some can move a considerable distance (up to 100μm in 2 weeks), there is no directional bias to this movement^13^. Since the net displacement of any cell is typically no more than 50μm with no direction bias and OPCs, maintain an evenly-spaced grid-like pattern despite developmental changes in their overall density and we assume that any OPCs that migrate into the ROI volume will be offset by an equal number of OPCs that migrate out^13^.
3) The total number of Olig2+ cells within the cortex and corpus callosum cannot be increased by the differentiation of cells that are not within the oligodendroglia lineage, cells that do not express the pan oligodendrocyte marker Olig2 or equivalents such as Sox10.

We intentionally sampled the septal component of the corpus callosum bound by the cingulate bundles laterally. In addition to ensuring anatomical landmarks that persist though development, there is good evidence that Sub Ependymal stem cell derived OPCs do not populate this region of the corpus callosum^30^. However, it is possible that a very small pool of SEZ-derived OPCs may be included as part of our analysis. SEZ-derived OPCs express PDGFRα and Olig2^30^, satisfying the assumption that the total pool of oligodendroglia can only be increased by cells committed to the oligodendrocyte lineage.

### Quantification and validation of cell death

To estimate the levels of cell death in the corpus callosum and cortex, we adopted a published protocol to identify pyknotic nuclei using Hoechst 33342 labelling^31^. For every age assessed, n = 3–4 and analysis was performed on high-resolution confocal images (Supplementary Figure 11). After P60, pyknotic nuclei were rare and it was not feasible to quantify these cells. In addition, the pyknotic nuclei counts were performed on individuals that were also subject to stereological analysis. This made it possible to identify the volumetric density of pyknotic nuclei at each age assessed (Figure 3C). To estimate total cell death in 24 hours per mm^3^ (Figure 3D), we used published data that cells in an irreversible stage of cell death remain in the tissue for ∼140 minutes prior to phagocytic clearance in the CNS^19, 32, 33^. Assuming that cells dye at random (i.e they are not all symmetrically dying at a single instance) it was possible to use our pyknotic nuclei density counts plus a cellular clearance time of 140 minutes^19^ to estimate daily cell death within a mm^3^ of either callosal or cortical parenchyma. To do this we used the following:

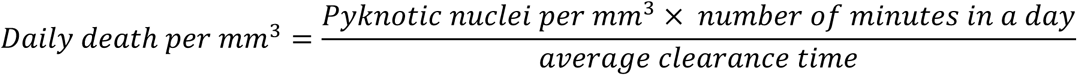

Where number of minutes in a day = 60 min x 24 = 1440;
average clearance time = 140 min.

Despite performing multilabel immunofluorescence (Supplementary Figure 11), it was not possible to assign a fraction of pyknotic nuclei specifically to the OL lineage. Therefore, the cell death values represent total cell death that occurs via pyknosis. Death in the OL lineage will account for some proportion of this total death. The % of pyknotic nuclei we identified was consistent with a previous assessment performed in the Optic nerve (Supplementary Figure 11), as well as with identified densities of activated caspase 3 cells in the cortex at similar ages^34, 35^. Assessing pyknosis is unlikely to capture 100% of all cell death. Cells may die by other means (eg. phagoptosis) and this would not be captured in our counts. It is likely that total cell death in vivo may exceed our estimates determined by counting pyknotic nuclei and modelling clearance times. Nevertheless, our data indicates that cell death within the OL lineage must account for a considerable proportion of the overall homeostatic levels of ongoing cell death that occurs in both cortical and callosal tissues in vivo.

### Sensitivity analysis

Our data revealed that OPCs have an average Tc of 35 h (Figure 1A). To provide additional validation, we modelled cumulative production in the corpus callosum as described previously except using a fixed OPC Tc of 35 hours. (Supplementary Figure 13). This generated a cumulative plot that was very similar to what is generated when we use measured values of Tc. We next asked the question; how sensitive is the system to a systematic over- or under-estimation of Tc or GF? To address this we systematically over- or underestimated Tc or GF and generated cumulative plots (Supplementary Figure 13). A systematic error in measuring either Tc or GF would lead to dramatic alterations in total cumulative production and such changes would be inconsistent with the levels of cell death we observed (Figure 3 and Supplementary Figure 11). Additionally, this analysis showed that the system is most sensitive to systematic changes in Tc: altering Tc by just one standard deviation of the measured value at each age had a profound impact on total cell production.

### Statistical software

All statistical tests performed are described in the relevant figure legends. We used GraphPad Prism version 8 for MacOS, GraphPad Software, San Diego, California USA, and where necessary, we validated non-linear regression modelling using Mathematica Version 11 (Wolfram Research).

## Supplementary Figures

**Supplementary Figure 1.**
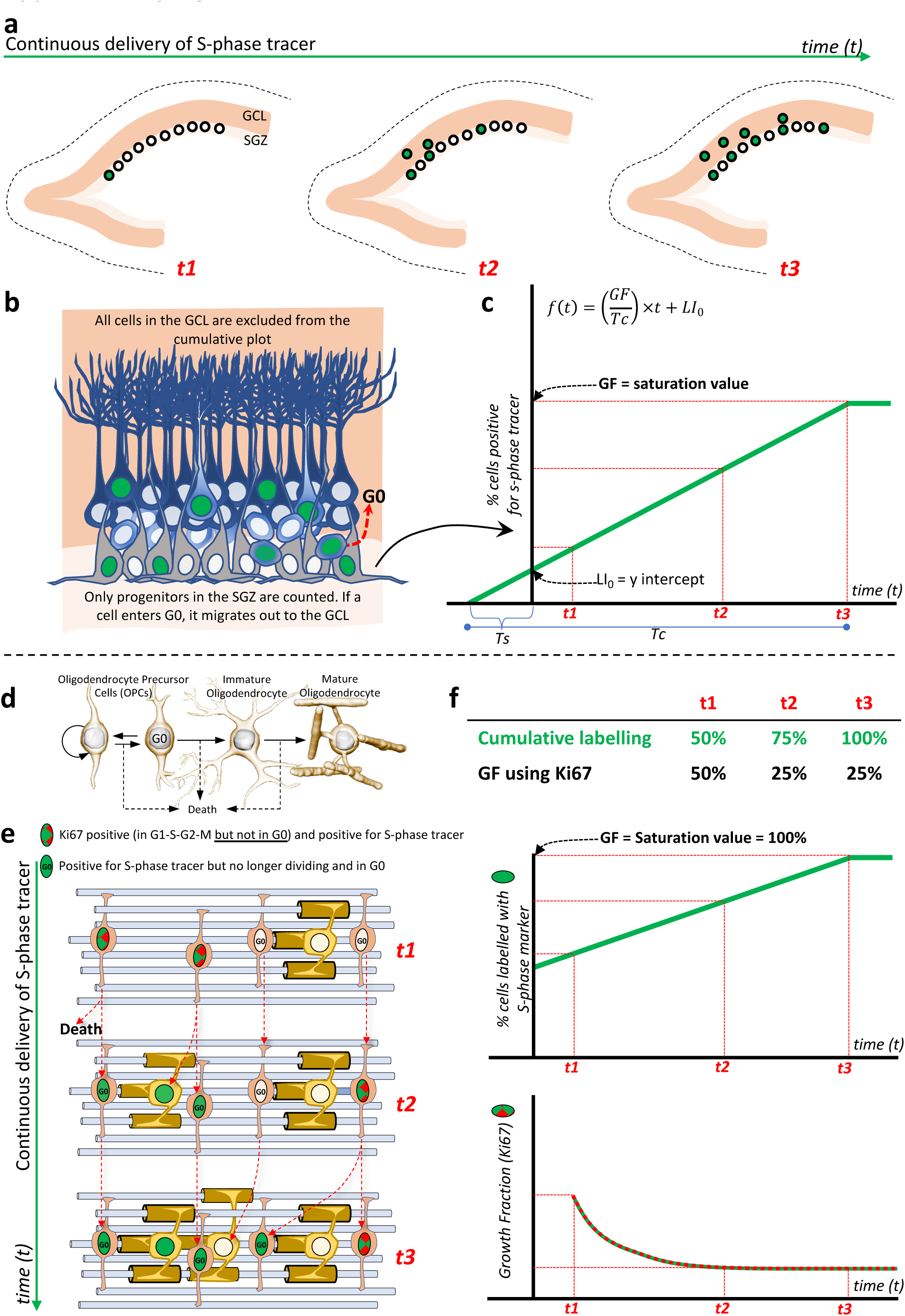
The cumulative labelling method: limitations and assumptions. A) Illustration of proliferating cells within the dentate gyrus of the hippocampus that incorporate the S-phase tracer into their DNA (labelled green) over time (t1, t2 & t3). Postmitotic cells retain the S-phase tracer, however they leave the anatomically defined proliferative subgranular zone (SGZ) and mitigate into the granule cell layer (GCL). B) The cumulative labelling method assumes that as soon as a cell enters G0 it migrates out of the anatomically-defined proliferative zone^3^, in this case the SGZ. The method also this assumes that no cells within the SGZ have the capacity to exit and re-enter the cell cycle, meaning the system is in a steady-state of growth^3^. C) Example of a cumulative labelling plot. When the biological assumptions are satisfied, plotting and determining the rate at which a population of cells becomes saturated makes it possible to determine key cell cycle parameters ^3^: The cell cycle length (Tc), the growth fraction (GF), the labelling index (LI0), and the S-phase length (Ts). The labels t1, t2 & t3 on the horizontal axis correspond to the same labels in the illustrations given in A). D) Image to illustrate that OPCs can divide, transition between dividing and non-dividing (enter and exit G0), differentiate, or die within the tissue. E) When an S-phase tracer is continuously delivered in this instance (t1, t2 & t3), dividing OPCs will incorporate it during S-phase (green). However, thymidine analogue S-phase tracers permanently label the daughters and there is no means to separate daughter OPCs that move in to G0 (labelled G0) from those activity dividing the system. Violation of this assumption leads to an overestimation of the GF^3^. To overcome this barrier, we used Ki67 to identify actively dividing cells ^10, 23^ (Supplementary Figure 3).

**Supplementary Figure 2:**
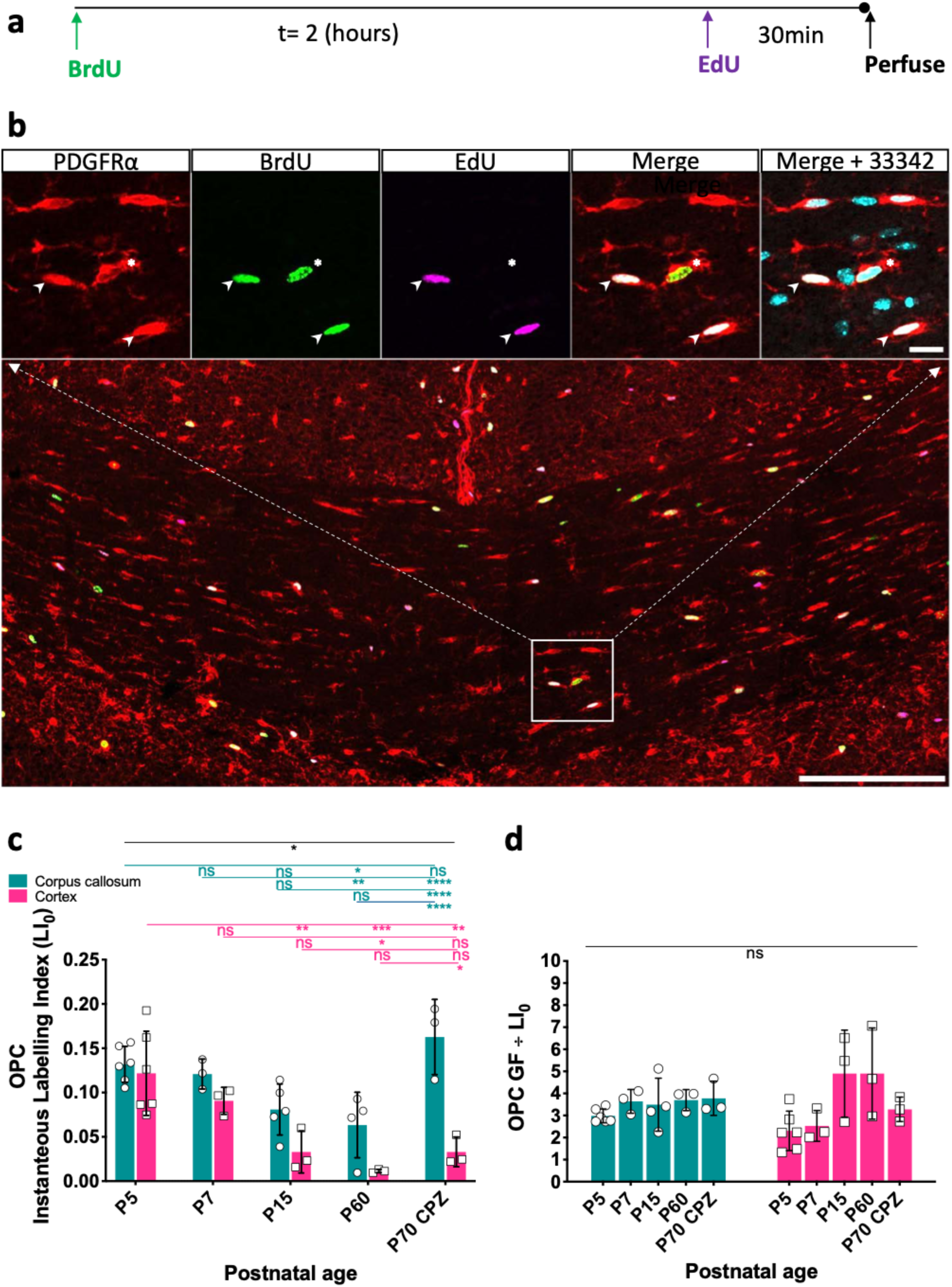
Double S-phase labelling and assessing the GF/LI0 ratio. **a)** Schematic of the double S-phase labelling method used. **b)** Illustration of double S-phase labelling at P5 in the corpus callosum. Arrow heads identify OPCs positive for PDGFRα, BrdU and EdU, demonstrating that these cells were in S-phase for the duration of the injection interval. Star identifies an OPC positive for PDGFRα and BrdU only, demonstrating a cell that transitioned though the S-phase prior to the injection of the second tracer (EdU). Scale bars in the lower and upper images indicate 200 µm and 10 µm respectively. **c)** The Instantaneous Labelling Index (LI_0_), is the fraction of cells in S-phase at any instant. This is significantly distinct between the corpus callosum and cortex and is elevated in response to 3 weeks of cuprizone demyelination. **d)** The ratio of GF/LI_0_ ratio does not significantly change across region, either with age, or in response to cuprizone demyelination. For all data; n=3–4 per data point. Statistics: 2-way ANOVA interaction (black line) and multiple comparisons (black connectors and coloured lines). Significance is indicated: ns = no significance and the p values: p<0.05, p<0.01, 0.001, and p<0.0001 represented by *, **, ***, **** respectively. Error Bars represent ± SD.

**Supplementary Figure 3:**
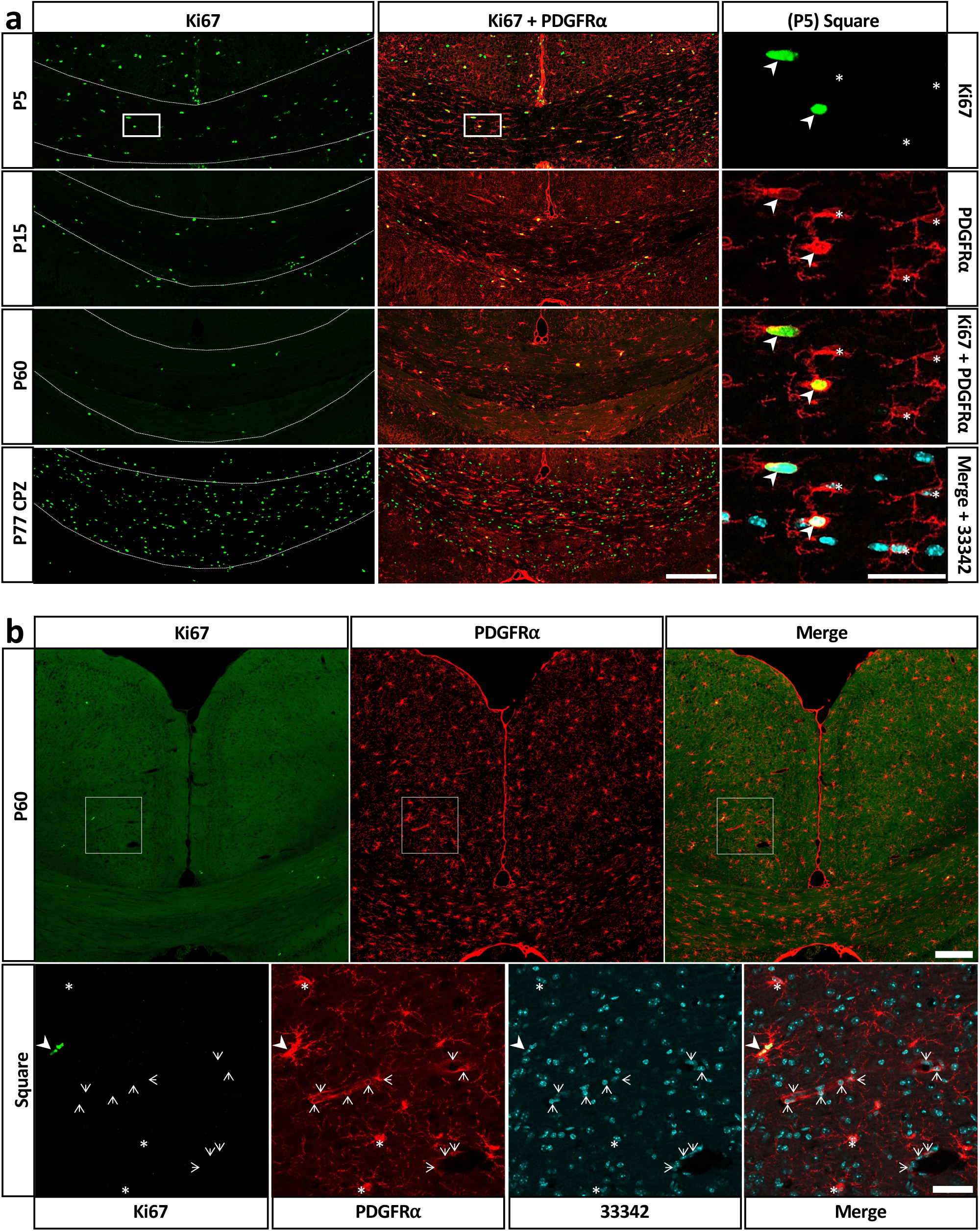
Assessing the GF using immunoreactivity for Ki67. a) The fraction of Ki67+ actively dividing OPCs decreases with age, however this is dramatically elevated in response to 3 weeks of cuprizone mediated demyelination (P77 CPZ). Arrow heads indicate actively dividing OPCs positive for both PDGFRα and Ki67. Stars denote OPCs positive for PDGFRα but not expressing Ki67. Bars: large image and small image 200 µm and 20 µm respectively. b) Image illustrating that in young adult mice (P60), few OPCs are positive for Ki67 under normal homeostasis. The enlarged square was selected due to the presence of blood vessels: it is possible to discriminate the morphological differences between PDGFRα+ OPCs (arrowheads and stars), and pericytes found lining blood vessels (open arrows)^27^. Care was taken to ensure PDGFRα positive pericytes were excluded from all analysis. Bars show 200 µm (large image) and 20 µm (small image).

**Supplementary Figure 4:**
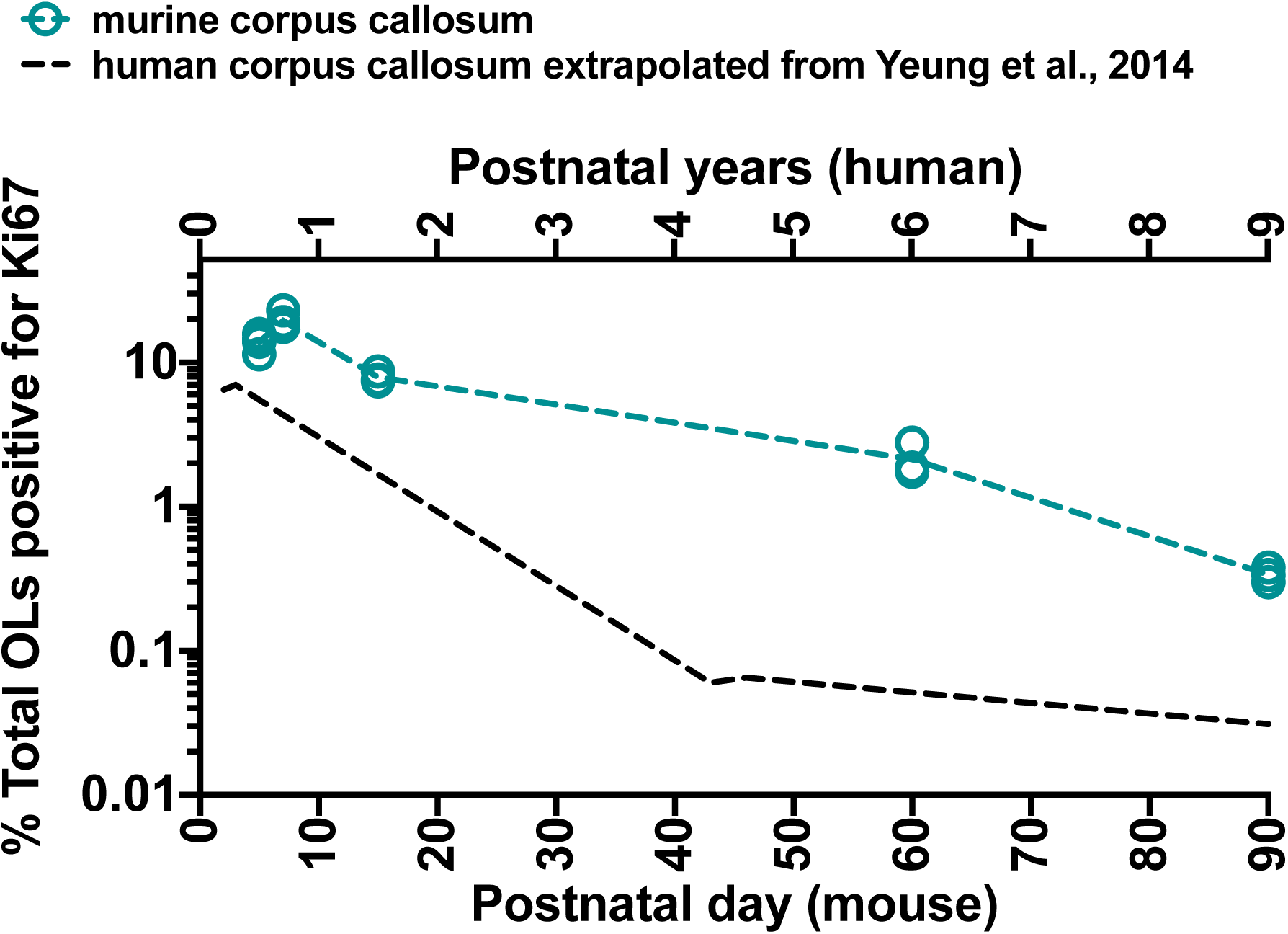
GF changes in the corpus callosum of the mouse and human. Representation of total oligodendroglia that are positive for Ki67 in the mouse (each open circle represents data from an individual animal where n=3-4/age. The total number of OPCs (PDGFRα+ Ki67+cells) represents a sub population of the total number of all oligodendroglia in the tissue. The data in this figure is distinct from the data represented in Figure 2C, in that here we express the Ki67 fraction in terms of all oligodendroglia as determined via stereology. This exercise is simply to compare the murine data we collected, against human data from Yeung et al., (2014) where the GF was represented as the fraction of all oligodendroglia that express Ki67. The dashed black line in this figure was generated by joining a straight line between values for individual subjects extrapolated from the article by Yeung et al 2014 (see Figure S4 G). We find that developmental changes in OPC GF occur in a similar way for both murine and human corpus callosum.

**Supplementary Figure 5:**
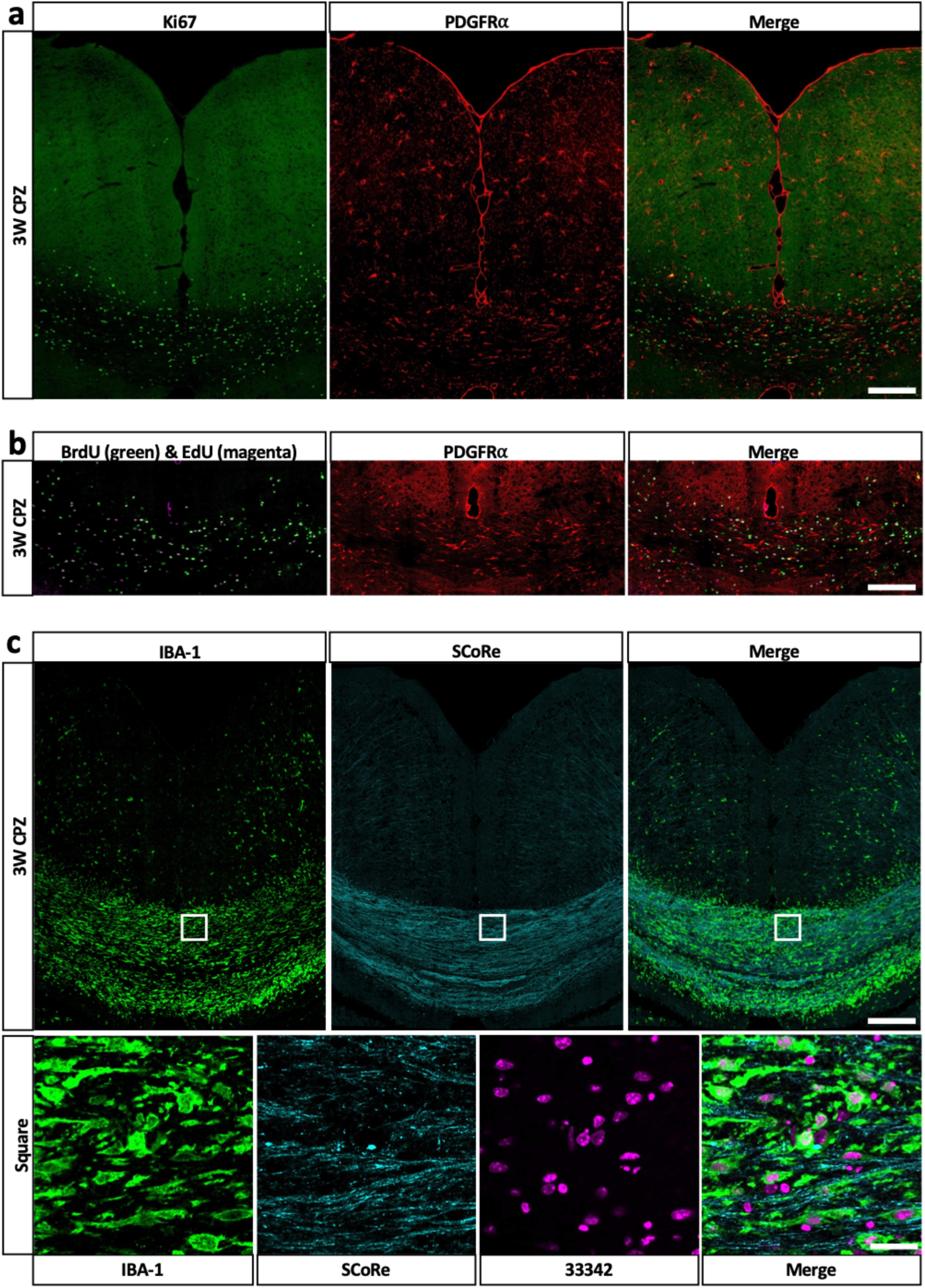
Cuprizone-induced OPC proliferation and demyelination. **a)** Ki67 staining in the corpus callosum is dramatically elevated in response to cuprizone demyelination. **b)** BrdU and EdU labelling within OPCs in the corpus callosum in response to cuprizone demyelination. **c)** Spectral Reflectance Confocal Microscopy (SCoRe) illustrating demyelination and IBA-1 macrophage activity in the corpus callosum resulting from 3 weeks of 0.2% cuprizone in normal chow. Error Bars 200 µm except for the enlarged square where the Bar is 25 µm.

**Supplementary Figure 6:**
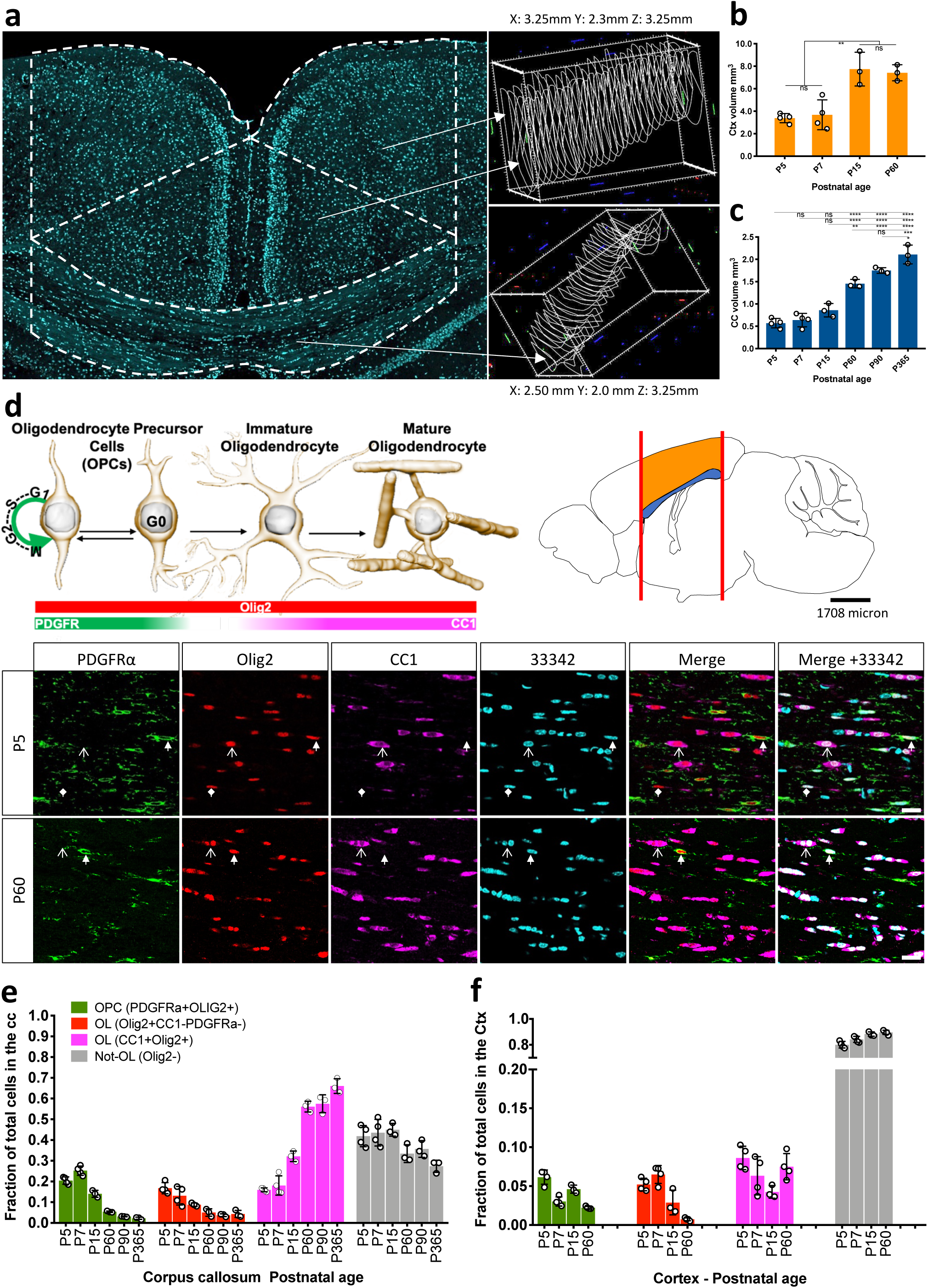
Stereology, volume assessments and phenotypic staining. **a)** Coronal section though a P60 mouse brain to illustrate how ROIs were captured for stereology. The cingulate bundles provided the anatomical landmark to limit the lateral margins of each trace, and the rostro-caudal margins were the midline unions of the corpus callosum: rostral, this was the most rostral aspect of the genu, and caudal, this was the most caudal aspect of the splenium. The white broken lines indicate a typical trace cortical and callosal trace. To the left of the image are collections of cortical and callosal traces from an individual at P60. The software used was stereoinvestigator V11 (MBF Bioscience). The dimensions for the bounding boxes surrounding the sets of traces are adjacent to the images. **b)** Total cortical volume increases significantly from P7 to P15. For all data: n=3–4 per age and the error bars represent ± SD. **c)** Total callosal volume increased significantly from P5 to P365. **d)** Antibodies directed against Olig2, PDGFRα and CC1 were used to identify stages within the oligodendroglial lineage. Oligodendroglia were identified as any cell positive for Olig2. OPCs were identified as cells positive for Olig2 & PDGFRα. Oligodendrocytes are cells positive for Olig2 & CC1, and cells positive only for Olig2 were considered immature oligodendrocytes that had lost the expression of PDGFRα, but yet to express CC1. **e)** Graphical representation of the proportions of cells in the corpus callosum, and **F)** cortex. For all data: n=3–4 per age; the error is ± SD; and the statistics in all ANOVA with Tukey’s post hoc testing significance is indicated: ns = no significance and stars; *, **, ***, **** represent p<0.05, p<0.01, 0.001, and p<0.0001, respectively.

**Supplementary Figure 7:**
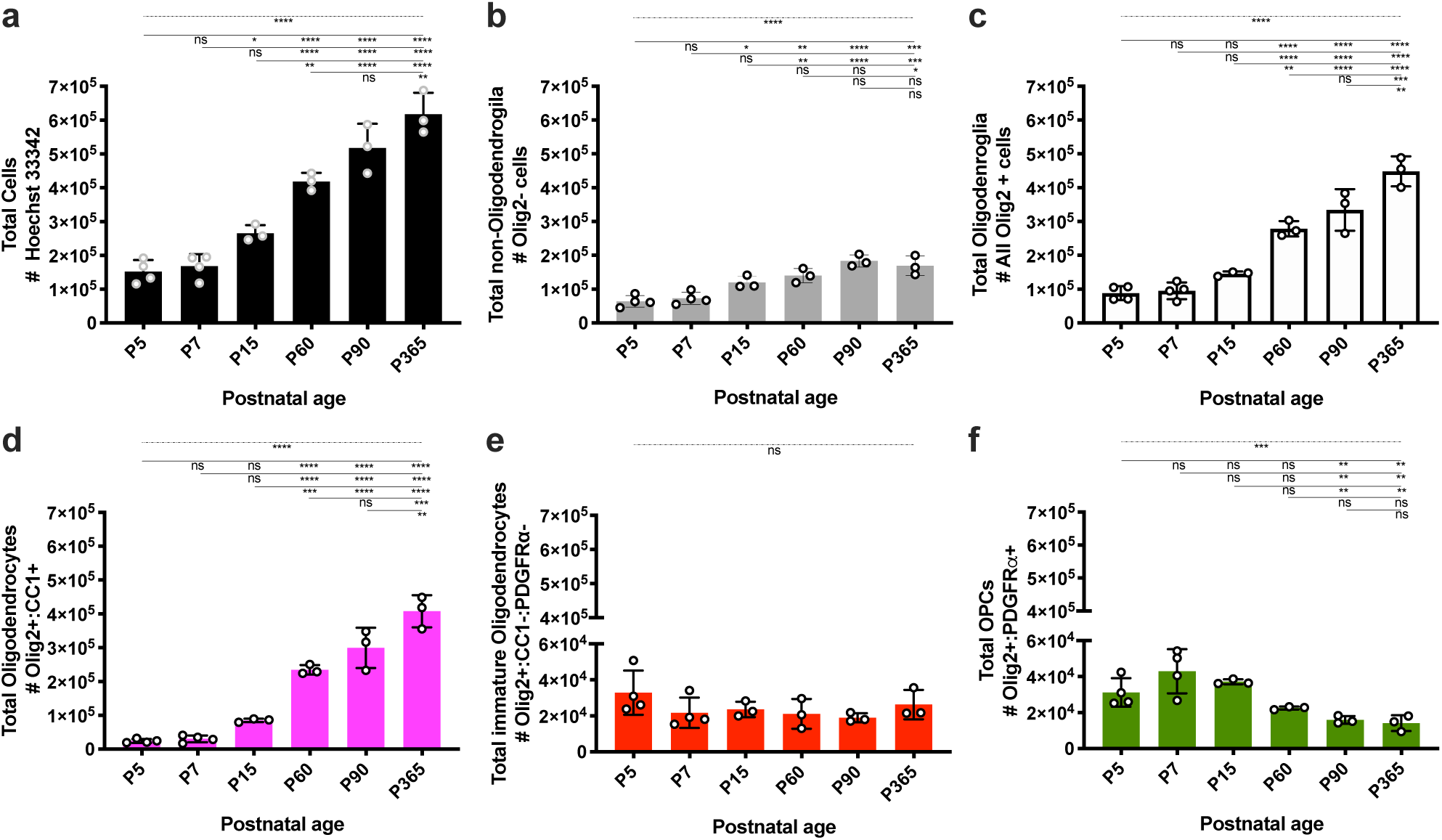
Change in total cell numbers in the corpus callosum. a) Hoechst 33342 was used to identify the total number of nuclei in the tissue. b) Total numbers of non-oligodendroglia (all cells positive for 33342 that lacked expression of the pan oligodendrocyte marker Olig2). c) Total number of oligodendroglia (all cells positive for 33342 and Olig2). d) Total number of oligodendrocytes (all cells positive for 3332, Olig2 and CC1). e) Total number of immature oligodendrocytes (all cells positive for 33342 and Olig2 that lacked expression of CC1 and PDGFRα). **a) f)** Total number of OPCs (all cells positive for 33342, Olig2 and PDGFRα). For all data: n=3–4 per age; the error is ± SD; and the statistics in all ANOVA with Tukey’s post hoc testing significance is indicated: ns = no significance and stars; *, **, ***, **** represent p<0.05, p<0.01, 0.001, and p<0.0001, respectively.

**Supplementary Figure 8:**
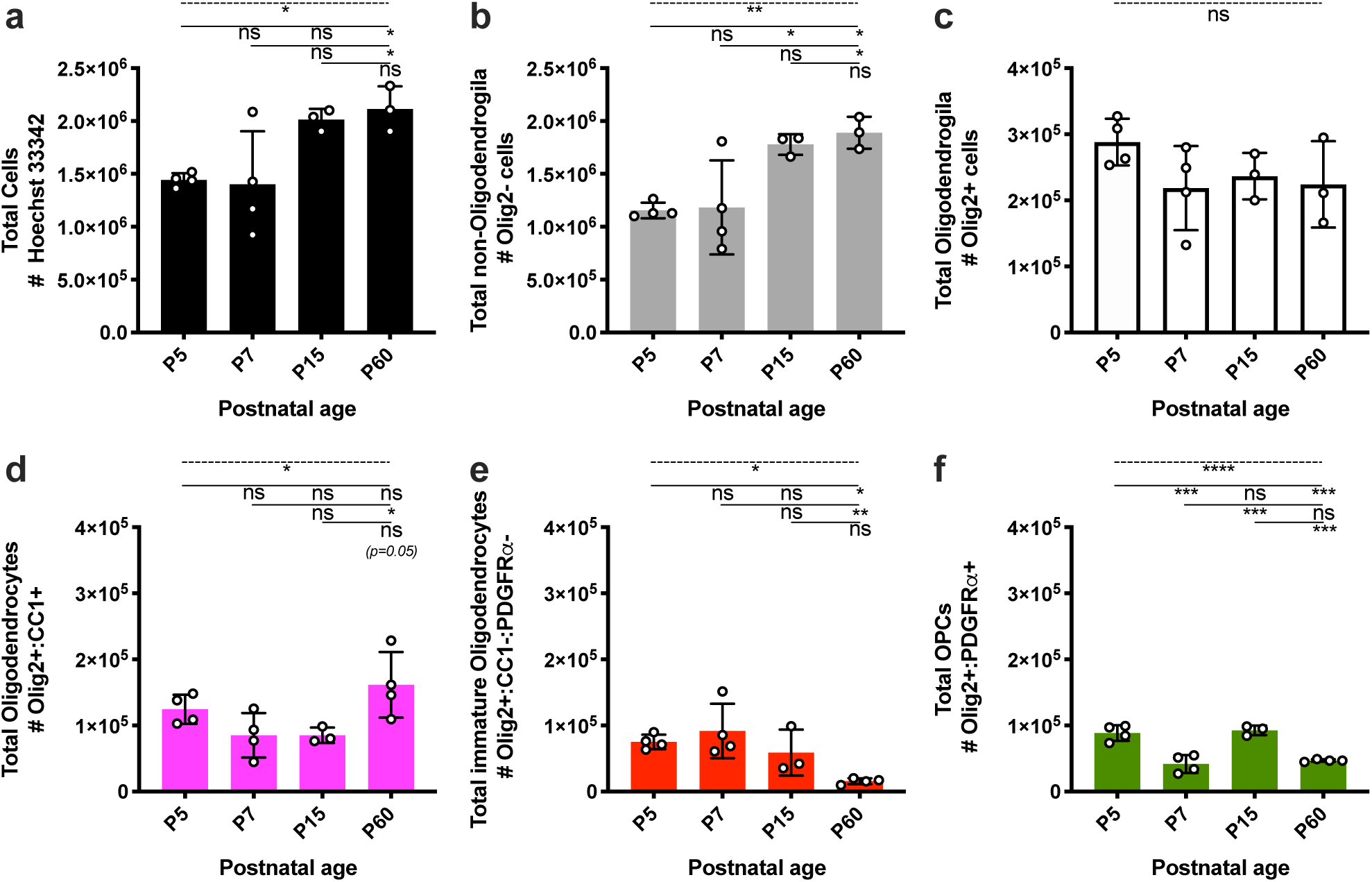
Change in total cell numbers in the cortex. **a)** Hoechst 33342 was used to identify the total number of nuclei in the tissue. **b)** Total numbers of non-oligodendroglia (all cells positive for 33342 that lacked expression of the pan oligodendrocyte marker Olig2). **c)** Total number of oligodendroglia (all cells positive for 33342 and Olig2). **d)** Total number of oligodendrocytes (all cells positive for 3332, Olig2 and CC1). **e)** Total number of immature oligodendrocytes (all cells positive for 33342 and Olig2 that lacked expression of CC1 and PDGFRα). **f)** Total number of OPCs (all cells positive for 33342, Olig2 and PDGFRα). For all data: n=3–4 per age; the error is ± SD; and the statistics in all ANOVA with Tukey’s post hoc testing significance is indicated: ns = no significance and stars; *, **, ***, **** represent p<0.05, p<0.01, 0.001, and p<0.0001, respectively.

**Supplementary Figure 9:**
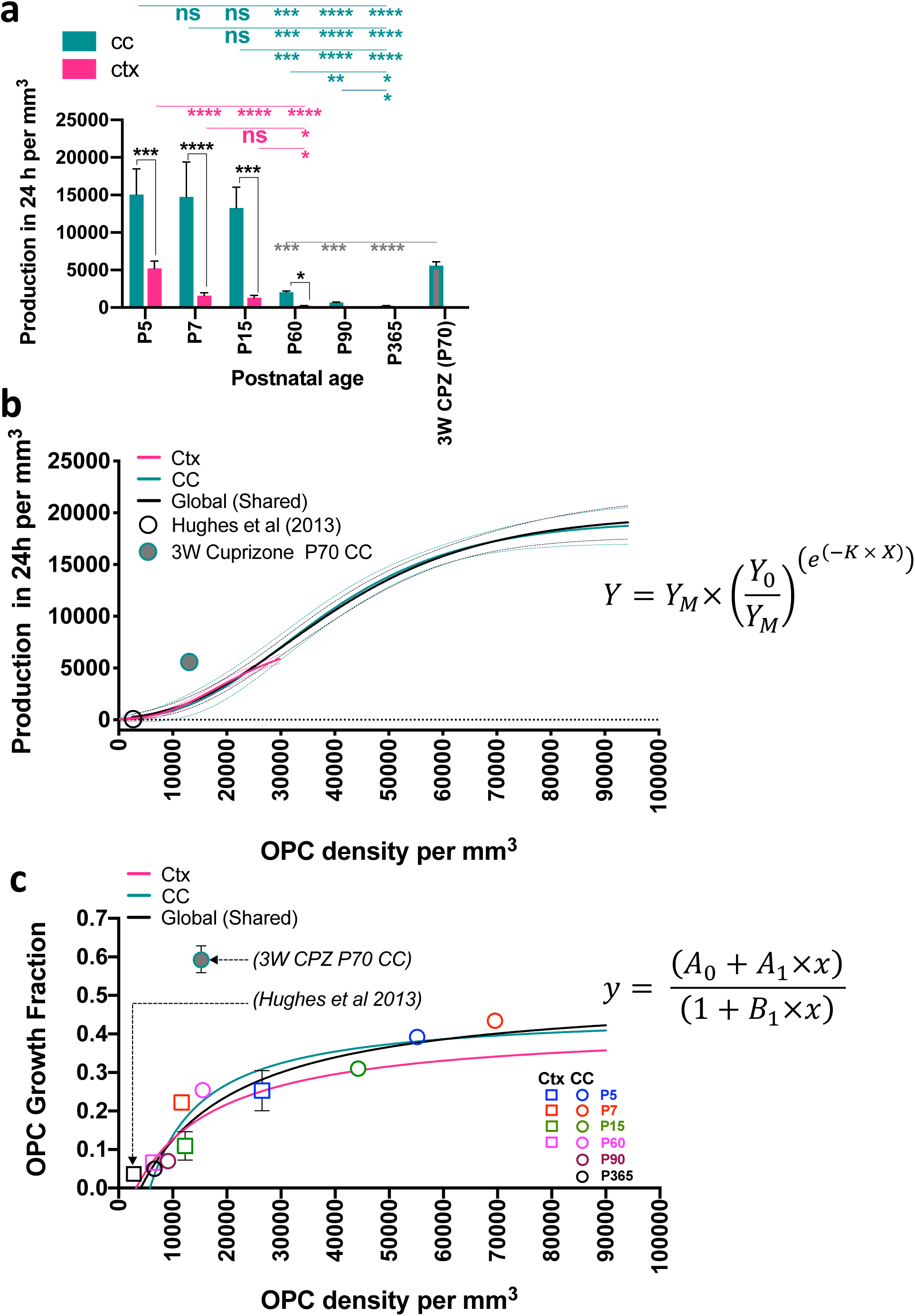
Positive relationship between OPC density, production and GF. **a)** Daily OPC production rates in the corpus callosum and cortex are strikingly distinct during development and after 3 weeks of cuprizone induced demyelination (3W CPZ P70). For each data point n= 3–5. Statistics: 2-way ANOVA analysis with multiple comparisons (black lines and stars), or 1-way ANOVA analysis (coloured lines and stars), significance: *, **, ***, **** = P<0.05, P<0.01, 0.001, and P<0.0001 respectively. Error bars = SD in all cases. **b)** Gompertz logistic regression analysis revealed that OPC volumetric density is positively related to their production rate (also see Figure 1D). A comparison of fits with the null hypothesis that one curve fits all data points was not possible to reject P = 0.5305. The values for all equations are listed below:

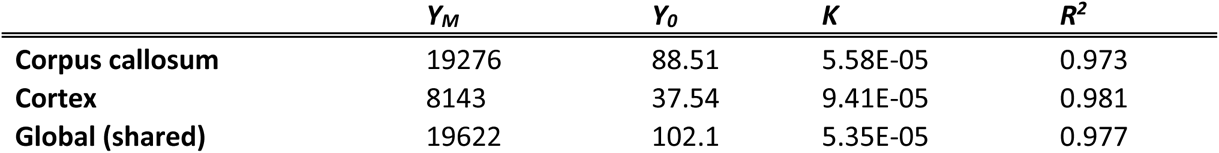 It was possible to compare our results to Hughes et al13, who report that cortical OPCs at a density of ∼160 cells per 0.06mm^3^ generate 0-7 cells per day (on average 3.5 cells per day) ^13^. We converted these values to mm^3^ and the plotted the data (black hollow circle) which fell within the 95% CI for all graphs (broken lines). Furthermore, if cortical OPCs are at a density of 160 cells per 0.06mm^3^ (or 2667 mm^3^), our cortex relationship predicts the daily number of OPCs generated to be 7 cells (SD ± 7 cells) per 0.06 mm^3^ per day. In addition to this, we plotted data derived from animals fed 3 weeks of cuprizone, this data point did not fall within the 95% CI intervals for any of the curves and indicates that OPCs production to density relationship is lost during injury. **c)** Padé (1,1) approximant (non-linear regression model) revealed OPC volumetric density is related to their GF. A comparison of fits with the null hypothesis that one curve fits all data points was not possible to reject P = 0.3715. The values are listed below:

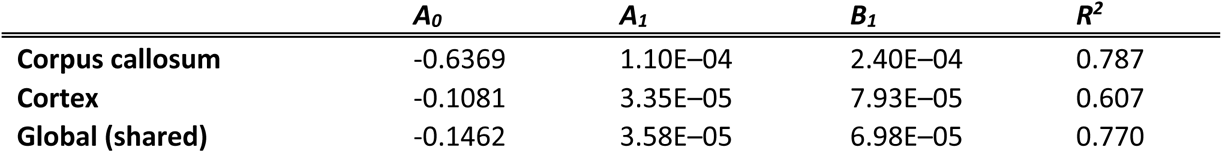 Using our data that OPC Tc is 35 h on average, along with the density and production rates identified by Hughes et al13, it was possible to estimate a GF for their cortical OPCs (black hollow Square):

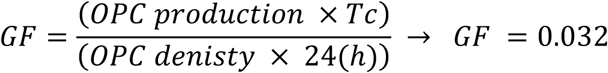 We plotted this GF value which fell very close to what was predicted by our cortical relationship. In addition to this, we plotted data derived from animals fed 3 weeks of cuprizone. The relationship between OPC production and density was strikingly distinct after injury. For each data point n= 3–5 and error bars = SD in all cases.

**Supplementary Figure 10:**
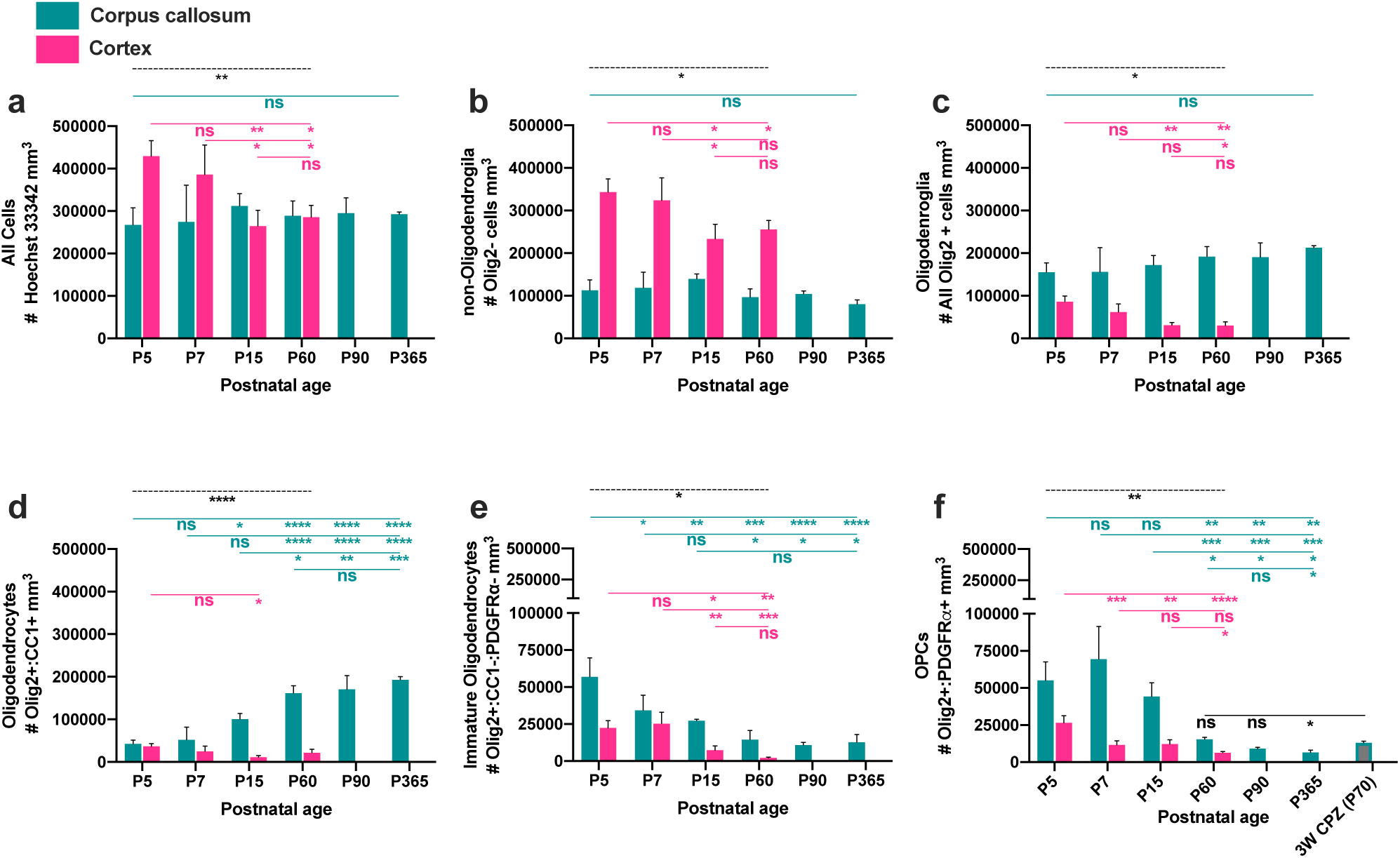
OL volumetric density in the corpus callosum and cortex. **a)** Hoechst 33342 was used to identify the density (mm^3^) of nuclei in the tissue. **b)** Total number of non-oligodendroglia per mm^3^ (all cells positive for 33342 that lacked expression of the pan oligodendrocyte marker Olig2). **c)** Total number of oligodendroglia per mm^3^ (all cells positive for 33342 and Olig2). **d)** Total number of oligodendrocytes per mm^3^ (all cells positive for 3332, Olig2 and CC1). **e)** Total number of immature oligodendrocytes per mm^3^ (all cells positive for 33342 and Olig2 that lacked expression of CC1 and PDGFRα). **f)** Total number of OPCs per mm^3^ (all cells positive for 33342, Olig2 and PDGFRα). For all data: n=3–4 per; the error is ± SD; and the statistics in all figures 2-way ANOVA testing (Indicated by black broken line) and ANOVA with Tukey’s post hoc multiple comparisons (Coloured lines). Significance is indicated, ns = no significance and stars; *, **, ***, **** represents p<0.05, p<0.01, 0.001, and p<0.0001 respectively.

**Supplementary Figure 11:**
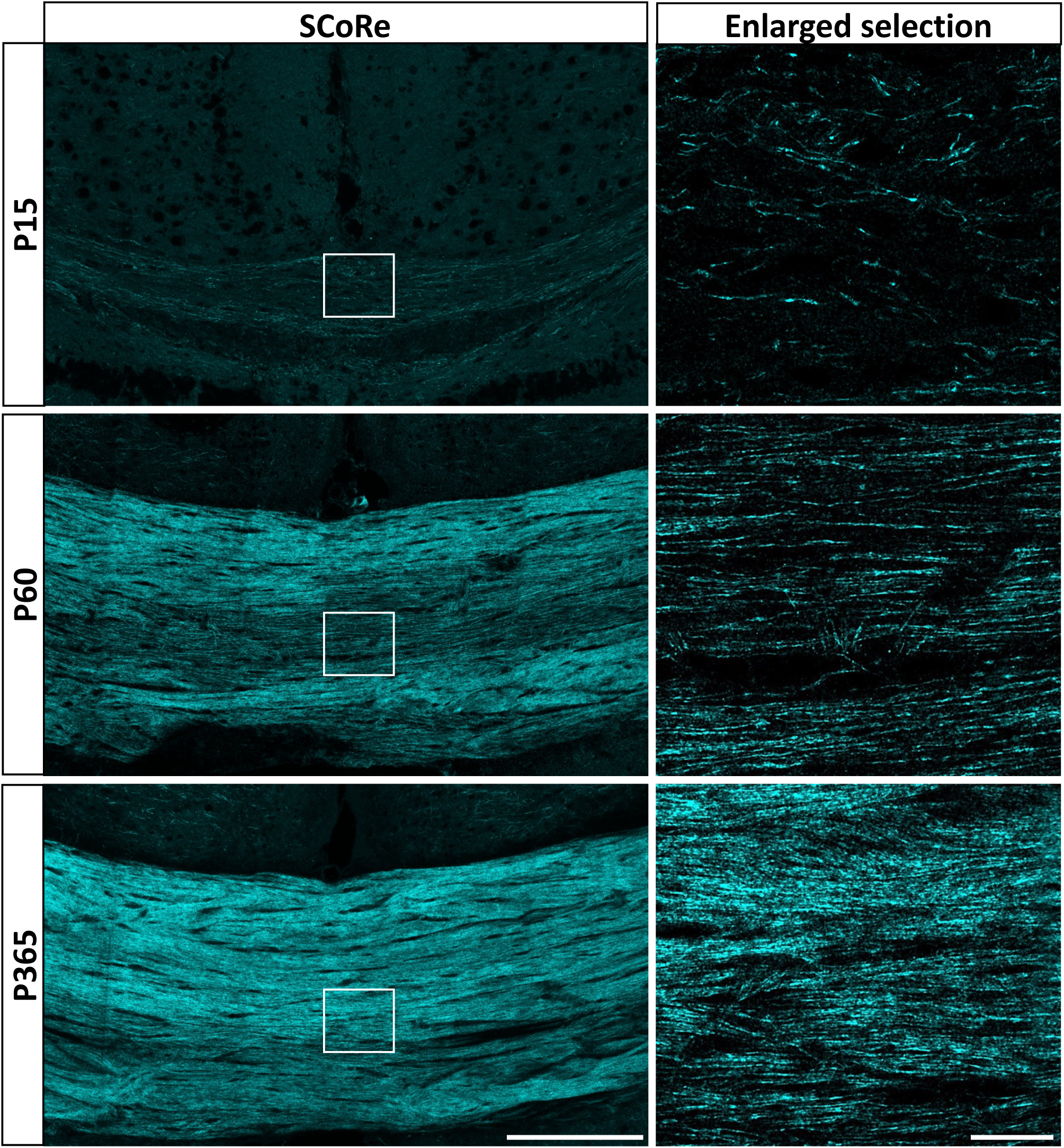
Age-related increase in myelin in the corpus callosum. SCoRe imaging of compact myelin in the corpus callosum, demonstrating an age-related increase in callosal area positive for SCoRE signal. Care was taken to ensure the coronal sections compared were from same relative anatomical position cutting from the Splenium of the corpus callosum forward. Large image bar = 200 µm, excerpt enlarged image bar = 20 µm.

**Supplementary Figure 12:**
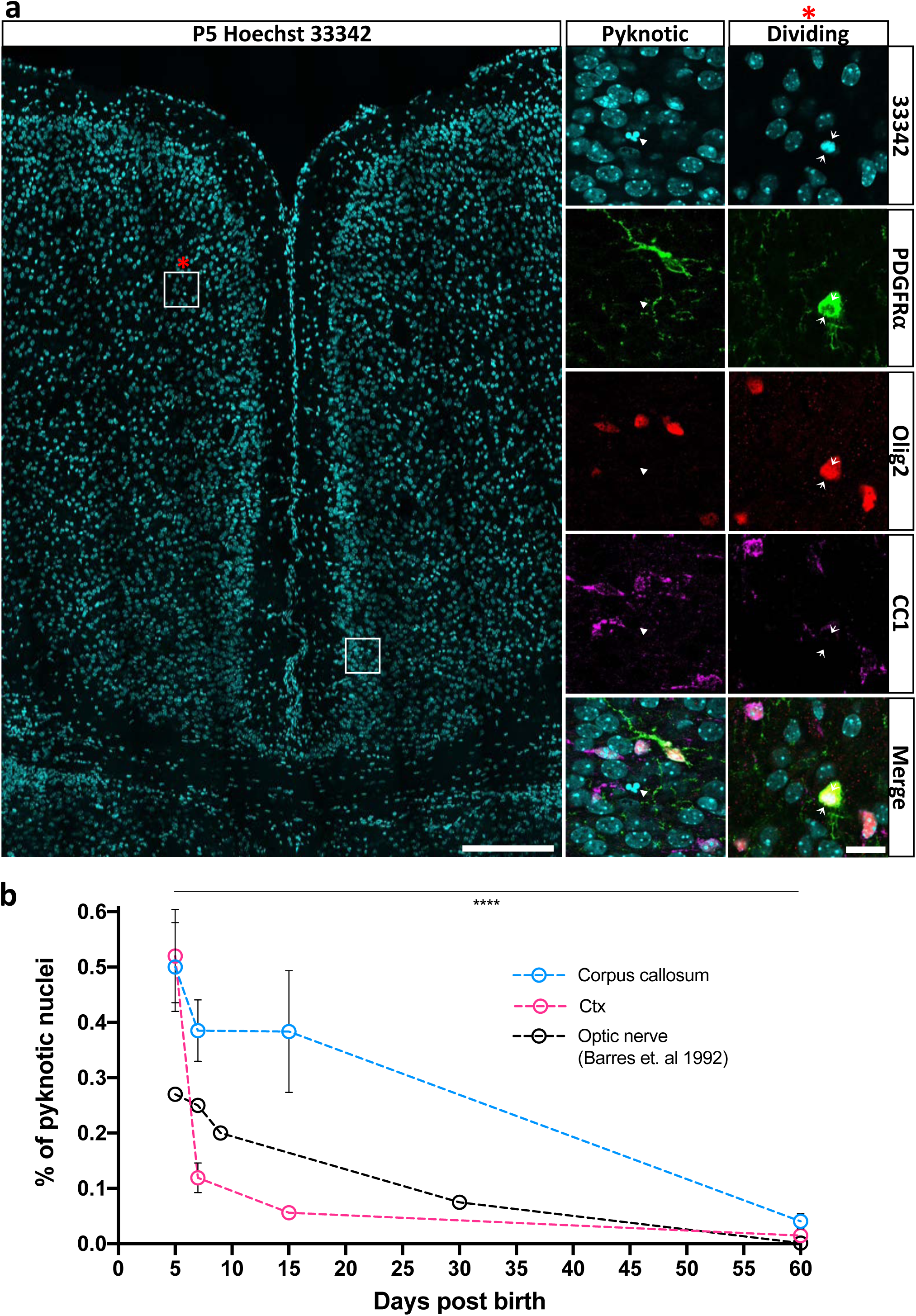
Identification and quantification of the % of pyknotic nuclei. **a)** Pyknotic nuclei were identified using Hoechst 33342^31^. Two ROIs are shown enlarged to show the pyknotic profile (the square without an asterisk) and the dividing nuclear profile (the square with an asterisk). Arrow heads indicate a pyknotic cell and paired arrows indicate the equatorial plane of a dividing OPC (positive for PDGFR⍺ and Olig2). Bar 200 µm and 10 µm. **b)** The % of all pyknotic nuclei in the corpus callosum and cortex respectively, as well as the fraction of pyknotic nuclei identified by Barres et al (1992) in the developing Optic Nerve^35^. Error bars ± SD. Statistics are from cell counts, n=3–4 for each data point, a 2-way ANOVA testing data from the corpus callosum and cortex, the significant interaction is indicated by stars**** = p<0.001.

**Supplementary Figure 13:**
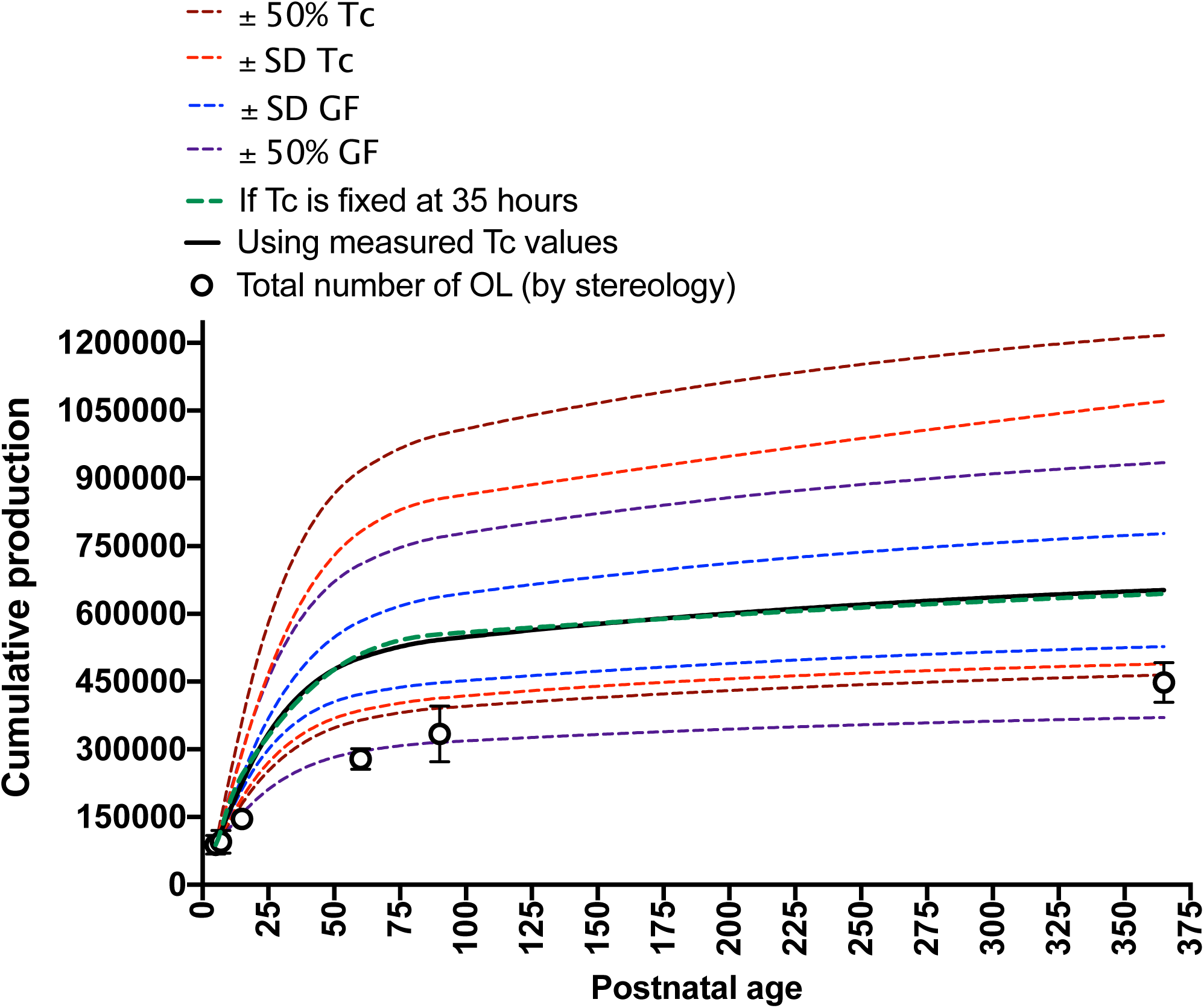
Sensitivity analysis: systematically varying Tc or GF dramatically alters cumulative production. To provide additional validation that OPCs Tc is stable over the long term, we modelled cumulative production in the corpus callosum using a fixed OPC Tc of 35 hours. This produced a similar curve (broken green line) to what was generated when the measured cell cycle values were used (solid black line). To illustrate how total cumulative production changes if GF, or Tc, values were systematically over or underestimated, we again modelled cumulative production with systematic alterations to only the GF or the Tc by: 1) a value equal to ± the standard deviation of the measured average value at each time point, or 2): ± 50% of the average measured value at each time point. Overlaid on the graph is the change in total OL number as determined by stereology (black circles). n = 3–4 for each measured data point and the error bars = ± SD.

## Acknowledgements

All images were collected at the Biological Optical Platform at the University of Melbourne, and the Florey Advanced Microscopy and Immunohistochemistry Service. In particular we would like to acknowledge Dr Carolina Chavez (FAMIS) for her help with stereoinvistigator; Dr Jessica L Fletcher and Dr David Homewood (Dept. Anatomy and Neuroscience University of Melbourne) for ongoing comments and input over the course of the project. This project was funded by: National Health and Medical Research Council (NHMRC) Early Career Fellowship (#GNT1111041 to DGG), Multiple Sclerosis Research Australia Betty Cuthbert Fellowship (#15-059 to DGG), NHMRC Project Grant (#GNT1058647 to JX), Australian research Council (ARC) Discovery Project Grant (#DP180102397 to JX) and ARC Discovery Project (#DP140100339 to BDH).

